# Differential modification of the C-terminal tails of different α-tubulins and their importance for microtubule function *in vivo*

**DOI:** 10.1101/2022.05.12.491705

**Authors:** Mengjing Bao, Ruth Dörig, Paula Vazquez-Pianzola, Dirk Beuchle, Beat Suter

## Abstract

Microtubules (MTs) are built from α-/β-tubulin dimers and used as tracks by kinesin and dynein motors to transport a variety of cargos, such as mRNAs, proteins, and organelles, within the cell. Tubulins are subjected to several post-translational modifications (PTMs). Glutamylation is one of them, and it is responsible for adding one or more glutamic acid residues as branched peptide chains to the C-terminal tails of both α- and β-tubulin. However, very little is known about the specific modifications found on the different tubulin isotypes *in vivo* and the role of these PTMs in MT transport and other cellular processes *in vivo*. In this study, we found that in *Drosophila* ovaries, glutamylation of α-tubulin isotypes occurred clearly on the C-terminal ends of αTub84B and αTub84D (αTub84B/D). In contrast, the ovarian α-tubulin, αTub67C, is not glutamylated. The C-terminal ends of αTub84B/D are glutamylated at several glutamyl side chains in various combinations. *Drosophila TTLL5* is required for the mono- and poly-glutamylation of ovarian αTub84B/D and with this for the proper localization of glutamylated microtubules. Similarly, proper Kinesin-1 distribution in the germline also depends on *TTLL5* as well as the refining of Staufen localization and the normal fast ooplasmic streaming with its directional movement, two processes known to depend on Kinesin-1 activities. In the nervous system, the pausing of anterograde axonal transport of mitochondria is affected by an enzymatically dead mutant of *TTLL5*. Our results demonstrate *in vivo* roles of *TTLL5* in differential glutamylation of α-tubulins and point to the *in vivo* importance of α-tubulin glutamylation for Kinesin-1-dependent processes.

**Summary:**

1. α-tubulin glutamylation was established in the C-terminal domain of *Drosophila* αTub84B and αTub84D (αTub84B/D). Multiple glutamyl residues were pinpointed in this domain. The female germline α-tubulin, αTub67C, is, however, not glutamylated.
2. *TTLL5* is required for mono- and poly-glutamylation of αTub84B/D.
3. *TTLL5* is required for the proper Kinesin heavy chain (Khc) distribution in the germline, and the kinesin-based refinement of Staufen localization and ooplasmic streaming during late oogenesis.
4. *TTLL5* is needed for the pausing of anterograde trafficking of mitochondria in axons.

## Introduction

Microtubules (MTs) are fundamental cytoskeletal filaments comprising heterodimers of α- and β-tubulins. Despite their high level of conservation in eukaryotes, MTs are still quite diverse because cells express different tubulin genes and proteins, which can undergo several different post-translational modifications (PTMs). Based mainly on *in vitro* assays, these variants are thought to optimize the MT interaction with different microtubule-associated proteins (MAPs) and motors.

*Drosophila melanogaster* possesses four α-tubulin genes. Three of them are expressed during oogenesis and early embryogenesis, suggesting that they contribute to the formation of microtubules during these stages. The three genes are *αTub84B*, *αTub84D,* and *αTub67C* (Kalfayan and Wensink, 1982). αTub84B and αTub84D differ in only two amino acids. Their primary sequence in the C-terminal domain is identical, and both are expressed in most tissues and throughout development. In contrast, αTub67C is expressed exclusively during oogenesis, where it is maternally loaded into the egg and embryo. The primary structure of αTub67C is distinctively different from αTub84B/D (Theurkauf et al., 1986). These differences are also apparent in the C-terminal domain and include the last residue, phenylalanine (Phe, F) in αTub67C and tyrosine (Tyr, Y) in αTub84B/D. *αTub85E* is the fourth α-tubulin. It is mainly expressed in the testes but not in the ovaries (Kalfayan and Wensink, 1982).

MTs are subjected to several PTMs, including the C-terminal tail glutamylation, glycylation, and detyrosination / tyrosination. Furthermore, acetylation and phosphorylation in the more central parts of the tubulins have also been identified (Arce et al., 1975; Eddé et al., 1990; Hallak et al., 1977; L’Hernault and Rosenbaum, 1985; Magiera and Janke, 2014; Redeker et al., 1994). Glutamylation is a PTM that adds one or more glutamic acids (Glu, E) to the side chains of glutamic acid residues in the C-terminal tails of both α- and β-tubulin, causing the formation of a branched peptide chain (Eddé et al., 1990; Redeker et al., 1992). Glycylation is another PTM that adds glycines to the γ-carboxyl group of glutamic acid residues in the C-terminal region (Redeker et al., 1994). At least in vertebrates, tubulin is also subjected to a particular cycle of de-tyrosination-tyrosination (Preston et al., 1979). In this case, the C-terminal Tyr or Phe of α-tubulin is cyclically removed, resulting in a C-terminal Glu residue, and a new Tyr can be re-added to the new C-term. The enzymes that catalyze glutamylation, glycylation, and tyrosination belong to the Tubulin-tyrosine-ligase-like (TTLL) family (Janke et al., 2005). In *Drosophila melanogaster*, this family encompasses 11 genes (flybase.org).

MTs have an intrinsic polarity with a plus and a minus end. On these tracks, kinesin motors transport cargo to the plus ends, and dynein motors move cargo toward the minus ends. Evidence for the importance of PTMs of tubulin subunits has already been reported *in vitro.* Artificial tethering of 10E peptides to the side chains at position E445 of α-tubulin and E435 of β-tubulin by maleimide chemistry increased the processivity (the run length) of kinesin motors on tubulin. For Kinesin-1, this was a 1.5-fold increase (Sirajuddin et al., 2014). In contrast to kinesin, Dynein/dynactin motors preferred a Tyr at the C-term of α-tubulin to initiate processive transport *in vitro* because the motor favored this isotype as an attachment point to dock onto the MTs (McKenney et al., 2016). Recent work now also demonstrated a role for the mouse tubulin glycylation enzymes. *In vitro* fertility assays showed that the lack of both *TTLL3* and *TTLL8* perturbed sperm motility, causing male subfertility in mice. Structural analyses further suggested that loss of glycylation perturbed the coordination of axonemal dynein, thus affecting the flagellar beat (Gadadhar et al., 2021).

*CG31108* is considered to be the homolog of the mammalian *TTLL5* gene and was therefore named *DmTTLL5 or TTLL5* (Devambez et al., 2017). It is required for α-tubulin glutamylation in the *Drosophila* nervous system (Devambez et al., 2017). The murine TTLL5 is composed of an N-terminal core tubulin tyrosination ligase-like (TTLL) domain, a cofactor interaction domain (CID), and a C-terminal receptor interaction domain (RID) (Lee et al., 2013). The N-terminal core domain was shown to provide the catalytic activity of TTLL5 and is highly conserved in *Drosophila* TTLL5 (van Dijk et al., 2007, Natarajan et al., 2017) (**Table 1**). According to FlyBase data, *Drosophila TTLL5* mRNA is expressed at high levels in ovaries (**Table S1**), even higher than in the nervous system where initial studies were performed (Devambez et al., 2017). We thus focused on *TTLL5*’s possible functions during *Drosophila* oogenesis to shed light on possible *in vivo* roles of glutamylated α-tubulins.

**Table 1.**
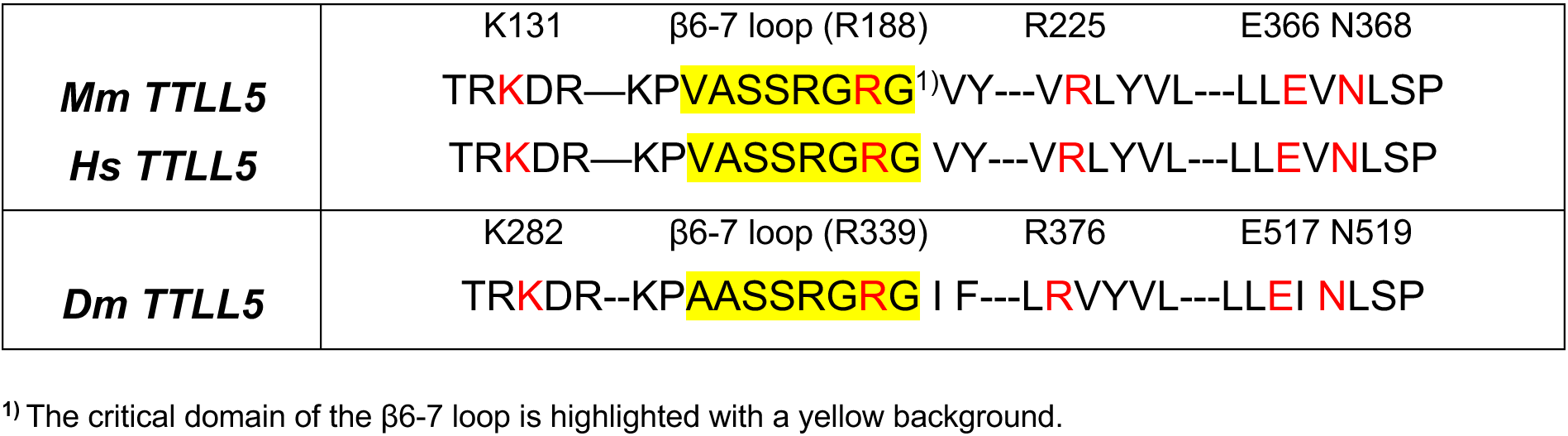
Conservation between mammalian TTLL5 and Drosophila TTLL5. Residues labeled in red are critical residues for α-tubulin glutamylation by murine TTLL5 and are conserved in human and Drosophila TTLL5 (van Dijk et al., 2007, Natarajan et al., 2017).

An *in vivo* mouse study showed that the absence of TTLL1-mediated polyglutamylation of α-tubulin increased the overall (both anterograde and retrograde) motility of mitochondrial transport in axons (Bodakuntla et al., 2021). However, another study in *Drosophila* showed that the vesicular axonal transport was not affected by the absence of TTLL5-mediated MT glutamylation in the segmental nerves of larvae (Devambez et al., 2017). Inspired by these controversial results, we also studied the function of *TTLL5* in a second tissue, the adult *Drosophila* wing nerves.

Our results revealed that *TTLL5* is essential for the normal glutamylation of the ovarian αTub84B/D. Surprisingly, even in wild-type ovaries, the oogenesis-specific aTub67C was not glutamylated, suggesting that this gene, *αTub67C,* may have evolved to produce MTs that contain regions that are less glutamylated. We found that *TTLL5* is required for the proper distribution of Kinesin heavy chain (Khc), for proper cytoplasmic streaming during late oogenesis as well as for fine-tuning the localization of the Staufen protein. Furthermore, *TTLL5* has a clear effect on the pausing of mitochondria during anterograde axonal transport in wing nerves. Our work, therefore, reveals the *in vivo* importance of glutamylation of specific α-tubulins.

## Results

### Specific glutamylation on multiple Glu residues of αTub84B/D, but not αTub67C in ovaries

Mass spectrometry was performed to analyze the glutamylation of the C-terminal tails of α-tubulins *in vivo*. Proteins extracted from ovaries were purified through SDS-PAGE, and α-tubulin bands were cut out and digested with the protease trypsin-N. C-terminal peptides of the different α-tubulin isotypes were then analyzed for post-translational modifications by liquid chromatography-mass spectrometry (LC-MS). Because the resulting C-terminal peptides are identical between the αTub84B and αTub84D, we refer to them as αTub84B/D (**Fig. 1A**). Analyses of wild-type ovarian extracts revealed that the general *Drosophila* α-tubulins αTub84B/D, but not the ovarian-specific αTub67C, showed glutamylation on their C-terminal tails (**Fig. 1A**). For αTub84B/D, side chain glutamylation was identified on all four glutamyl residues in the tail region, Glu443, Glu445, Glu448, and Glu449. The most extended side chains identified were 3Es in wild-type ovaries **(Fig. 1A,1B)**. According to FlyBase, *αTub84B+D* mRNAs are 4.7x as highly expressed as *αTub67C*. We identified a similar ratio (4.6x) of C-terminal peptides by the MS analysis (**Table S2**). Because of the lower abundance of Tub67C C-terminal peptides, we cannot rule out that rare glutamylation on the maternal αTub67C exists. Altogether, we conclude that glutamylation of α-tubulin had a strong preference for the C-terminal domain of αTub84B/D.

**Fig. 1.**
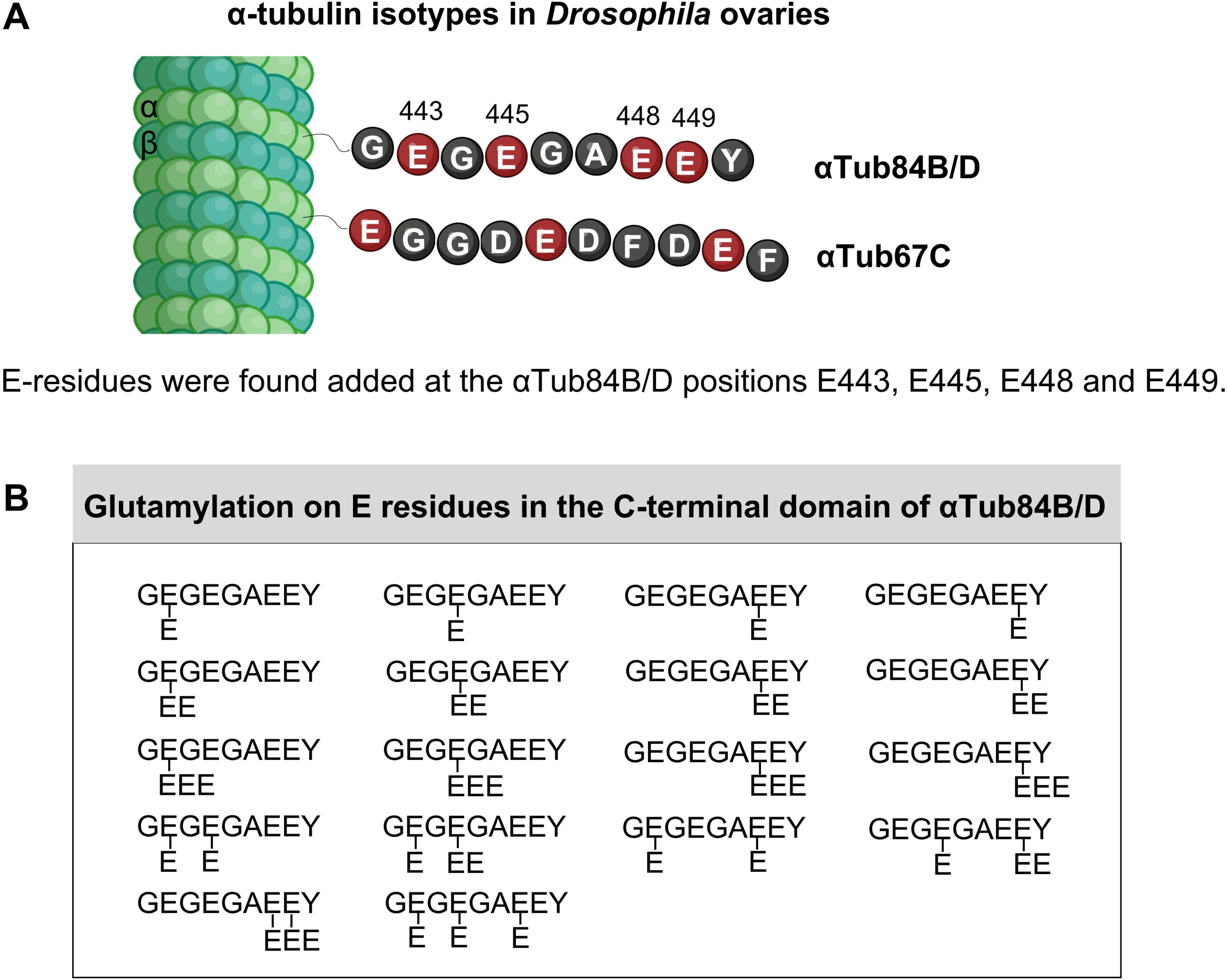
Mass spectrometry analysis of glutamylation of the C-terminal domains of *Drosophila* α-tubulins. A) Schematic representation showing how the C-terminal regions of αTub84B/D can be modified by glutamic acid residues added to the side chains of E443, E445, E448, and E449 in *Drosophila* ovaries. B) The C-terminal peptides containing the complete primary sequence were analyzed. The structure of the different C-terminal tail peptides of αTub84B/D modified by glutamylation is shown starting with G442. Different combinations of Glu side chain modifications were found added at positions E443, E445, E448, and E449 of αTub84B/D. Schemes were drawn by biorender.com.

### *Drosophila TTLL5* is required for MT mono- and polyglutamylation of ovarian αTub84B/D

To study the function of TTLL5, distinct *TTLL5* loss-of-function mutations were created and characterized (**Fig. 2A**). The *TTLL5^pBac^* and *TTLL5^Mi^* mutant strains carry a piggyBac and a Minos transposon, respectively, within the open reading frame of *TTLL5*. The *TTLL5^MiEx^* mutant was generated by imprecise excision of the Minos element, which caused the formation of a premature stop codon at gene position 8,132 in the open reading frame (codon position 392). An anti-TTLL5 antibody that cross-reacts with additional proteins in ovarian extracts also recognized a band corresponding to the calculated mass of TTLL5. This band was absent in ovarian extracts prepared from the three *TTLL5* mutants, suggesting that they are protein nulls for *TTLL5* or make only a truncated protein (**Fig. 2B**). A transgene containing the wild-type *Drosophila TTLL5* sequence fused to the Venus fluorescent protein sequence under UAS control (UASP-*Venus::TTLL5*) was also produced as a tool to rescue the phenotypes produced by the *TTLL5* mutations and for overexpression studies (**Fig. 2A**).

**Fig. 2.**
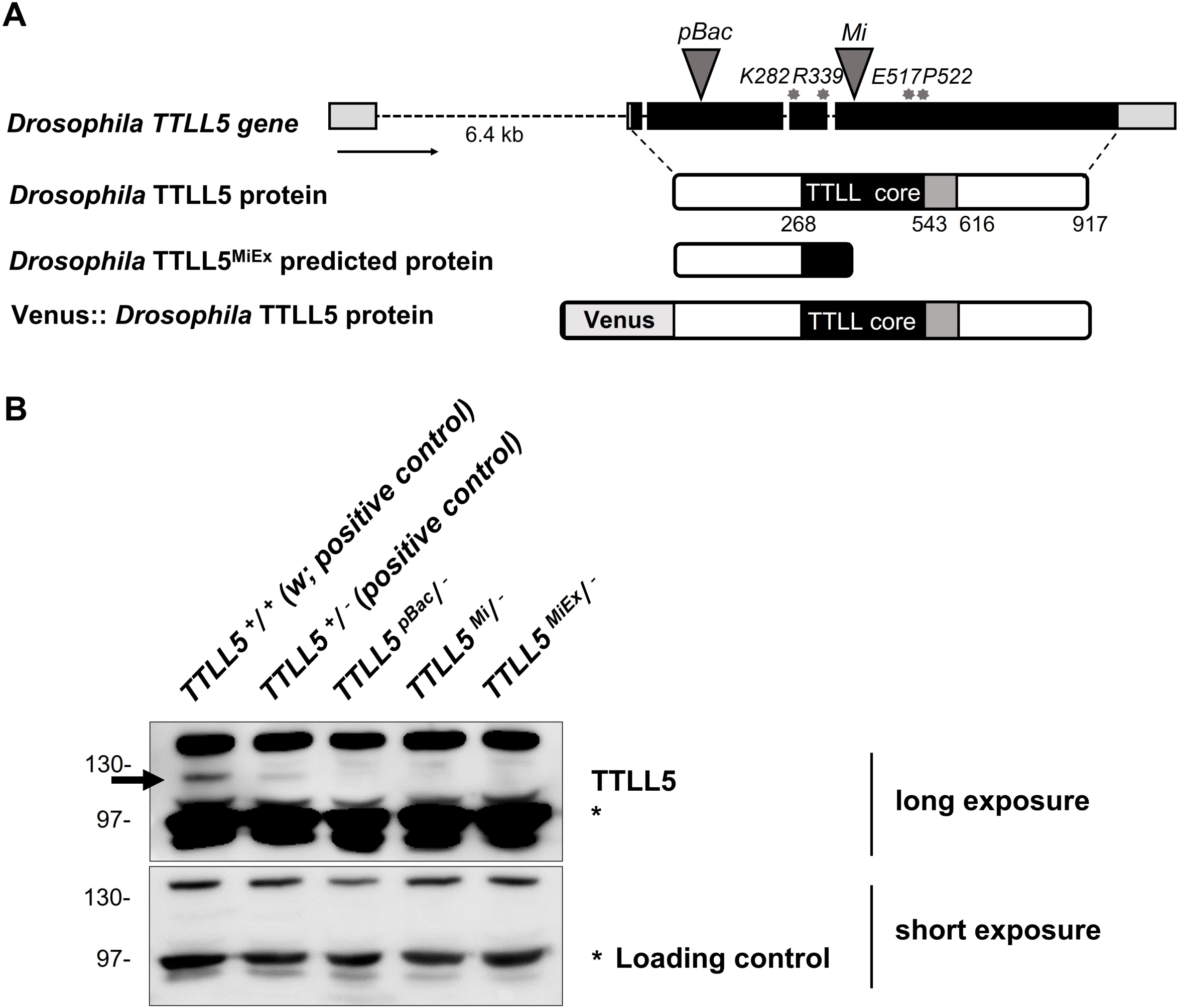
Schematic representation of the *Drosophila TTLL5* gene, *TTLL5^pBac^*, *TTLL5^MiEx^*, and Venus::TTLL5 proteins. A) The *TTLL5* gene structure is based on the FlyBase data for the *CG31108-RA* transcript. The positions of the loss-of-function mutations are marked on the *TTLL5* gene. Grey triangles: transposon insertions. Grey asterisks: codons targeted for generating InDels and point mutations. All *TTLL5* alleles were analyzed as hemizygous animals over *Df(3R)BSC679*, which removes the region of the *TTLL5* gene. B) Western blot from the soluble fraction of ovarian extracts. *w* (*TTLL5^+^/^+^),* which contains two wild-type copies of *TTLL5*, expressed the highest levels of TTLL5. The TTLL5 signal is strongly reduced or abolished in the null mutants *TTLL5^pBac^/^-^, TTLL5^Mi^/^-,^* and *TTLL5^MiEx^/^-^* compared to *w.* The hemizygous *TTLL5^+^* ovaries (*TTLL5^+^/^-^*) expressed less TTLL5 than *w.* The signal of an unspecific band (labeled with *), produced after a short exposure, was used as the loading control. In the following, we will refer to the genotypes as *w* for the control (*TTLL5^+^/^+^*) and *TTLL5^allele^* for the hemizygous mutants *(TTLL5^allele^/ ^-^*).

To quantify the effect of *TTLL5* on the glutamylation of the C-terminal tails of α-tubulins, we determined the frequency of C-terminal peptides containing the entire primary sequence and 0, 1, 2, or 3 Glu (E) modifications. This analysis was performed with extracts from wild-type ovaries and the different *TTLL5* mutant ovaries. Wild-type ovaries revealed that 62% of the peptides were modified, 41% with 1E, 14% with 2Es, and 7% with 3Es **(Table 2)**. On the other hand, no glutamylation was observed in *TTLL5^pBac^* and *TTLL5^MiEx^* mutants. This clearly shows that *Drosophila TTLL5* is essential for the side chain glutamylation of the C-terminal Glu443, Glu445, Glu448, and Glu449 of αTub84B/D and it appears to affect already the addition of the first Glu.

**Table 2.**
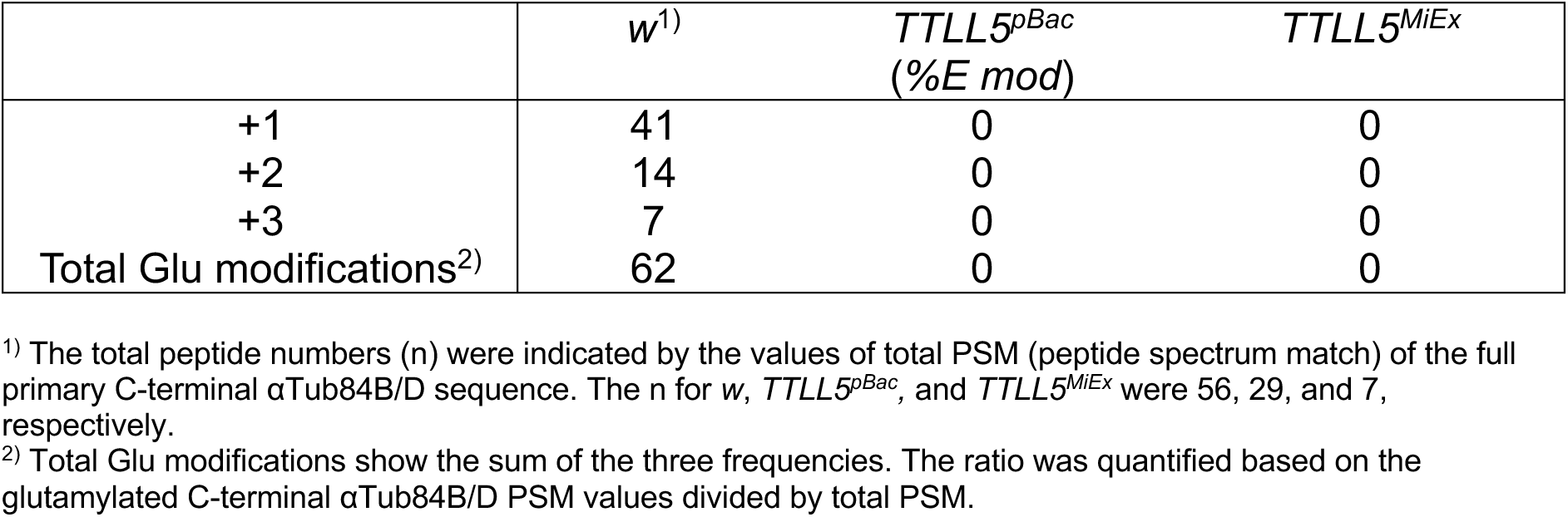
Role of TTLL5 for the glutamylation of the C-terminal domain of αTub84B/D. Frequency of glutamylated C-terminal peptides (containing the entire primary sequence) in w controls, TTLL5^pBac,^ and TTLL5^MiEx^ mutants containing modifications of 1E, 2E, or 3Es.

|  | *w*<sup>1)</sup> | TTLL5<sup>pBac</sup><br>(%E mod) | TTLL5<sup>MiEx</sup> |
| --- | --- | --- | --- |
| +1 | 41 | 0 | 0 |
| +2 | 14 | 0 | 0 |
| +3 | 7 | 0 | 0 |
| Total Glu modifications<sup>2)</sup> | 62 | 0 | 0 |

<sup>1)</sup> The total peptide numbers (n) were indicated by the values of total PSM (peptide spectrum match) of the full primary C-terminal αTub84B/D sequence. The n for *w*, TTLL5<sup>pBac</sup>, and TTLL5<sup>MiEx</sup> were 56, 29, and 7, respectively.
<sup>2)</sup> Total Glu modifications show the sum of the three frequencies. The ratio was quantified based on the glutamylated C-terminal αTub84B/D PSM values divided by total PSM.

We also measured the levels of monoglutamylation of α-tubulin by Western blotting using the GT335 antibody. This antibody reacts with the first glutamate on the glutamate side-chain even if it is polyglutamylated (**Fig. 3A**) (Wolff et al., 1992). Note that the GT335 antibody does not reliably detect all mono glutamylated residues in *Drosophila* tubulins because GT335 preferentially recognizes the first Glu attached to a Glu residue in an acidic environment. In mice, the middle Glu in the sequence -G**E**E- of the C-terminal tail of α-tubulin is such an “acidic” site. Once glutamylated, the branched Glu modification can be recognized by GT335 (Bodakuntla et al., 2021). In contrast, in *Drosophila* α-tubulin, both E443 and E445 in the sequence -G**E**G**E**G- are flanked by Glycines on both sides, which does not provide the acidic environment needed for efficient recognition by GT335. The GT335 signal may, therefore, mainly stem from the glutamylation on the consecutive Glu residues E448 and E449 in the sequence -GA**EE**Y. On Western blots from wild-type extracts, the GT335 antibody produced several bands. The band pointed out by the arrowheads is at the expected position, is reduced or absent in the mutants but reappears in the rescued ovary extracts (**Fig. 3B**). This result, therefore, supports the finding that *TTLL5* is required for monoglutamylation of α-tubulin, and it shows that the antibody signal is specific and can be used to monitor *Drosophila TTLL5* activity.

**Fig. 3.**
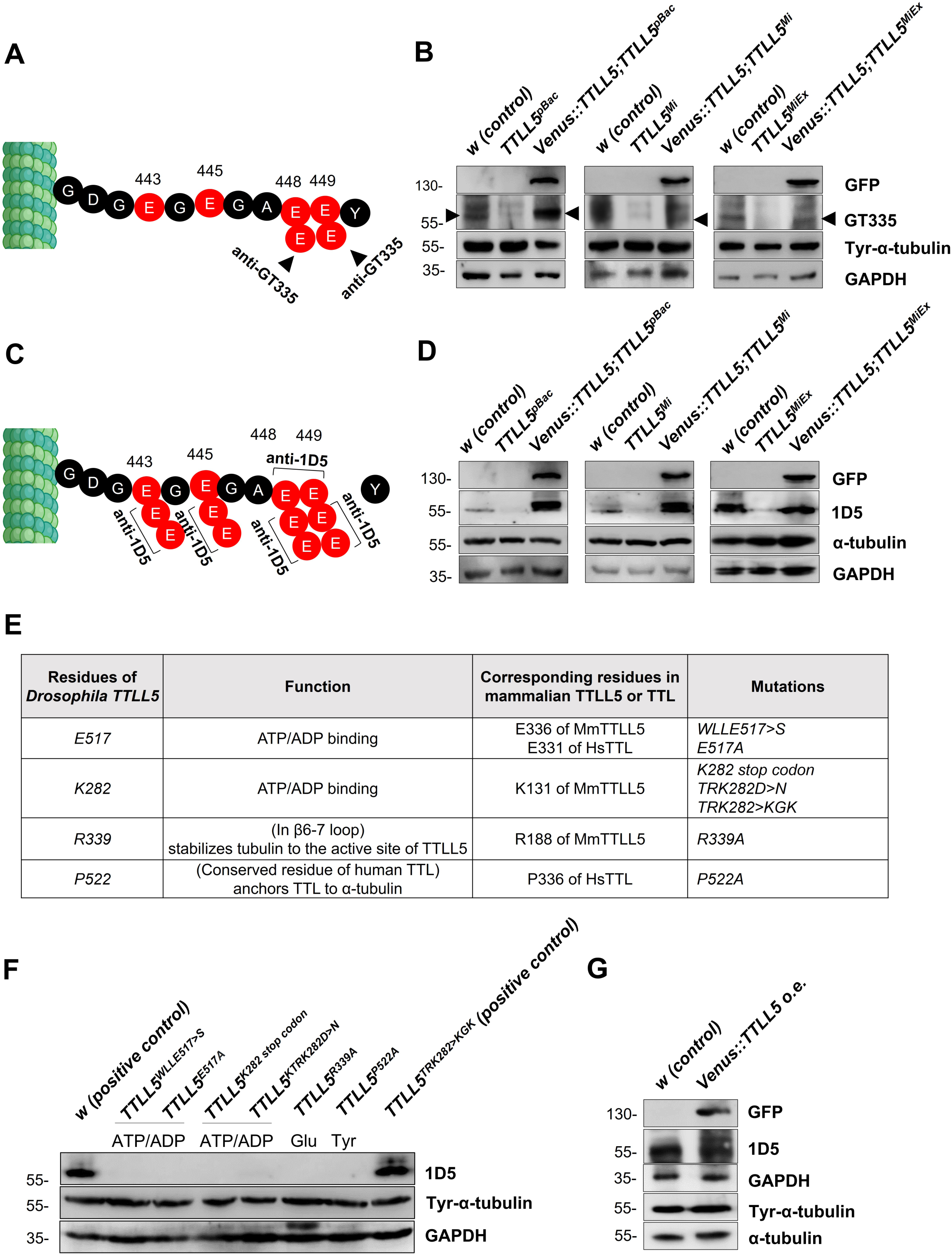
*TTLL5* is required for the glutamylation of α-tubulin in ovaries. A), C) Sites on αTub84B/D expected to be recognized by the antibody against monoglutamylated α-Tubulin (GT335; A) and by the 1D5 antibody recognizing polyglutamylated αTub (C). B), D) The glutamylation signal, produced by Western blotting with the GT335 and 1D5 antibodies, respectively, is lower in *TTLL5^pBac^, TTLL5^Mi^*, and *TTLL5^MiEx^* than in the *w* control but is restored and even elevated in mutant ovaries that express *Venus::TTLL5* under *MattubGal4* control. The up-shifted bands recognized by 1D5 can be seen in the rescued *TTLL5^pBac^* and *TTLL5^Mi^* mutants. The total tyrosinated α-tubulin levels seemed unaffected by the TTLL5 levels. α-tubulin and GAPDH served as loading controls. E) The genotypes and rationale of the *TTLL5* alleles generated by CRISPR/Cas9 are listed. The selected residues are predicted to be either important for the glutamylation or tyrosination function based on the alignment shown in **Tables 3** and **4**. F) Western blotting shows that the polyglutamylation signal is absent in all point mutants. *w* with two copies of *TTLL5 ^+^* is the wild-type control and the hemizygous *TTLL5^TRK282>KGK282^* still contains the crucial K282 residue and behaves like the control. Relative to the loading controls, 1D5 signals decrease slightly in the hemizygous situation with only one *TTLL5^+^* copy. G) Stronger up-shifted bands were observed with the polyglutamylation antibody (1D5) upon *Venus::TTLL5* overexpression in wild-type ovaries. Total α-tubulin levels were similar in samples with excessive expression of *TTLL5.* GAPDH was a loading control. All mutant *TTLL5* alleles were analyzed as hemizygous animals over *Df(3R)BSC679*. The genotypes of the rescued animals were *MattubGal4>UAS-Venus::TTLL5;TTLL5^alleles^/Df(3R)BSC679.* The genotype of the *Venus::TTLL5* overexpressing animals was *MattubGal4>UAS-Venus::TTLL5*.

**Table 3.**
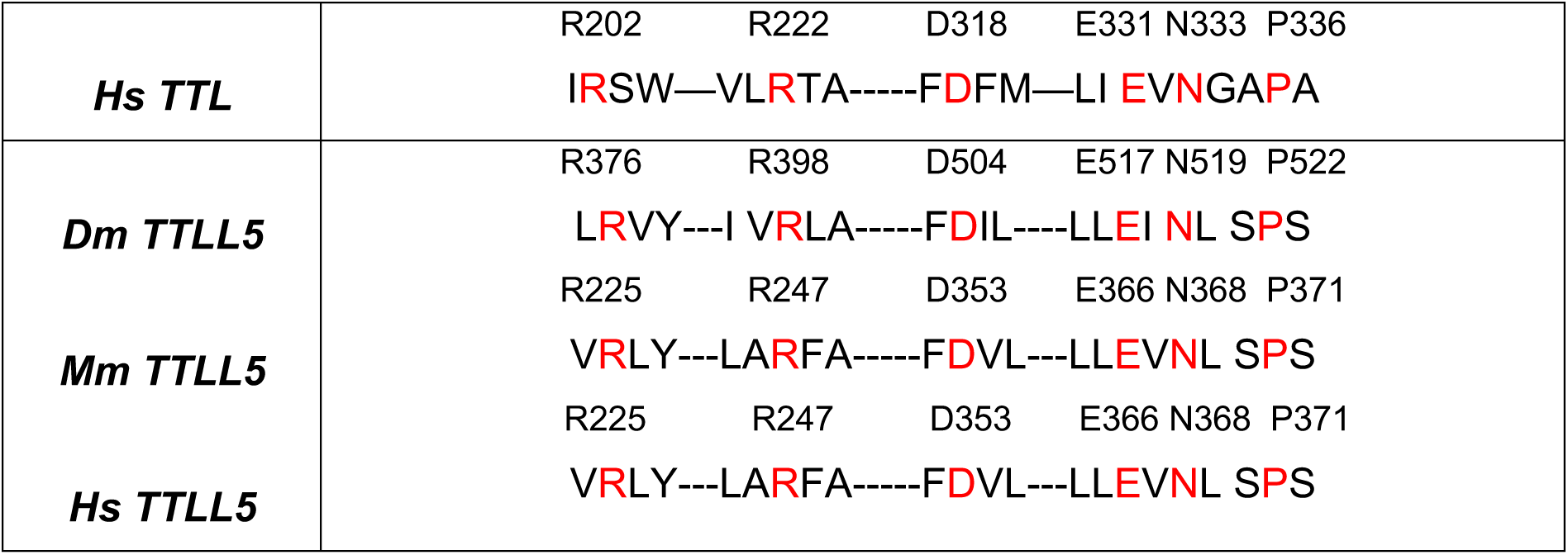
Conservation between TTL and TTLL5. Many critical **r**esidues of human TTL (red) are conserved in *Drosophila,* mouse, and human TTLL5 (Prota et al., 2013).

The 1D5 antibody recognizes α-tubulin with an -XEE sequence at the C-terminal α-carboxylate group (with X being a variable residue) (Rüdiger et al., 1999) **(Fig. 3C)**. Therefore, in principle, all Glu residues E443, E445, E448, E449, and the C-terminal Glu residues E448/449 can be recognized if they are not followed by another amino acid (such as Tyr) at their α-carboxylate group. Because *w* control ovaries showed high 1D5 cross-reactivity with a polypeptide of the predicted size, this, therefore, indicates the presence of glutamate side chains containing at least two glutamates and possibly C-terminally de-tyrosinated α-tubulin (Gadadhar et al., 2017; Preston et al., 1979) **(Fig. 3D)**. Consistent with the MS analysis, the *TTLL5* mutants showed a strongly reduced signal with 1D5, indicating that loss of *TTLL5* function prevents normal polyglutamylation. The 1D5 signal was again restored in all *TTLL5* mutants upon expression of the rescue construct. Overexpression of TTLL5 in *TTLL5^pBac^* and *TTLL5^Mi^* even produced a stronger 1D5 signal and an up-shifted band, suggesting that α-tubulin gets hyperglutamylated **(Fig. 3D)**. These results reveal that *TTLL5* is essential for the glutamylation of the α-tubulin subunit. Moreover, the result that the rescue construct induced the appearance of hyperglutamylated α-tubulin also suggests that, in normal cases, the endogenous *TTLL5* levels are usually limiting in ovaries.

Most of the murine TTLL5 residues involved in α-tubulin glutamylation (Natarajan et al., 2017) are conserved in *Drosophila* TTLL5 **(Table 1;** Natarajan et al., 2017). Two are known to be essential for the general enzymatic activity of TTLLs, and two for the interaction with the C-terminal domain of α-tubulin **(Fig. 3E)**. To determine whether these residues are indeed needed *in vivo* for the enzymatic activity of *Drosophila TTLL5* towards αTub84B/D, we used CRISPR/Cas9 to mutate the conserved codons in *Drosophila TTLL5. Drosophila* TTLL5 residues K282 and E517 correspond to mouse K131 and E336, respectively. They directly interact with ADP/ATP and thus affect general TTLL5 enzymatic activity (Natarajan et al., 2017). TTLL5^R339^ corresponds to mouse TTLL5^R188^, which is vital for TTLL5’s glutamylation activity. MmTTLL5^R188^ resides in the β6-7 loop, which forms a salt-bridge interaction with the C-terminal tail of α-tubulin and orients it towards the ATP/ADP-binding site (van Dijk et al., 2007). Glutamylation of α-tubulin was impaired in all these *TTLL5* mutants where the conserved residue was replaced **(Fig. 3F)**. Since these mutations did not abolish the stability of the corresponding mutant protein (**Fig. S1**), our results revealed the importance of E517, K282, and R399 of *Drosophila* TTLL5 for glutamylation. Most active site residues of human TTL (Prota et al., 2013; van Dijk et al., 2007) are also conserved between mammalian and *Drosophila* TTLL5 **(Table 3)**. We also mutated the *Drosophila TTLL5^P522^* codon corresponding to human *TTL^P336^* (and human *TTLL5^P371^*/mouse *TTLL5^P371^*), which is required for anchoring TTL to α-tubulin by forming hydrogen bonds with C-terminal tail residues of α-tubulin (Prota et al., 2013). However, the importance of *TTLL5^P371^* for glutamylation has not been reported in mammalian systems. Surprisingly, P522 turned out to be essential for the glutamylation of tubulins in *Drosophila* ovaries, revealing P522 as a novel critical residue for the glutamylation function of TTLL5. We also tested whether the overexpression of *TTLL5* in a wild-type background affected polyglutamylation similarly. Indeed, *UASP-Venus::TTLL5* overexpression in ovaries led to hyperglutamylation, as evident from the stronger signal and the additional upshifted band **(Fig. 3G)**. All these results point to *TTLL5’s* essential role in the glutamylation of α-tubulin.

### Other PTMs of the C-terminal tail of ovarian α-tubulin

Western blotting showed no or only minor changes in tyrosinated α-tubulin levels in any of the null or CRISPR/Cas9 generated point mutants, and *TTLL5* overexpression did also not affect these levels **(Fig. 3B, 3F, 3G)**. TTLL5 levels, therefore, do not appear to affect levels of tyrosinated α-tubulin in ovaries.

Our MS results also detected signals that could correspond to glycylated C-terminal α-tubulin peptides in ovarian samples. Such signals were observed in αTub84B/D but not in αTub67C. However, the standard procedure for the MS analysis involves the alkylation of Cysteine residues in the sample and this procedure can also modify Glu residues, giving rise to the exact same mass change as glycylation (+57.021464 Da; Kim et al., 2016). Our MS analysis could, therefore, not distinguish if a particular Glu residue was artificially modified by carbamidomethylation or naturally by glycylation. To independently test for this PTM, we also performed Western blotting experiments with ovarian extracts and probed them with the anti-mono Gly antibody TAP952 (Bré et al., 1996). Because *TTLL4A* is also expressed in ovaries **(Table S1)**, we initially considered it as a potential candidate for performing tubulin glycylation and included mutant samples of *TTLL4A* and *TTLL5* in this analysis. However, even in the wild-type control, the anti-mono Gly antibody TAP952 did not reveal a clear signal at the predicted position (**Fig. S2A**), suggesting that α-tubulin is not glycylated in *Drosophila* ovaries. Furthermore, because the known *Drosophila* polyglycylases (TTLL3A/B) were shown to be the main glycylases in the whole fly and are not expressed in ovaries (Rogowski et al., 2009)(**Table S1**), at present, the combined evidence does not warrant further analysis of this potential PTM in *Drosophila* ovaries.

To test for other possible functions of *TTLL4A* in ovaries, ovarian extracts from *TTLL4A* mutants were also blotted and probed with the anti-mono- and polyglutamylation antibodies GT335 and 1D5, respectively. The results showed a strongly reduced signal of both GT335 and 1D5 in the ovaries of the *TTLL4A^pBac^* mutant, and a weaker reduction of GT335 and 1D5 in the *TTLL4A^pEPgy^* mutant (**Fig. S2B, S2C**), suggesting the *Drosophila* TTLL4A might function as a glutamylase on α-tubulin in flies. We will discuss the possible interpretations of these results in the Discussion section.

### Presence of polyglutamylated microtubules in ovaries

We then analyzed the distribution of polyglutamylated microtubules during oogenesis by immunofluorescence using the 1D5 antibody. Specific 1D5 immunofluorescence signals were observed in the follicle cells and oocytes throughout the previtellogenic, middle, and late oogenesis stages (**Fig. 4**). In the stage 10B (S10B) oocyte, the 1D5 signal showed a slightly biased localization in the cortical region of the oocyte. We observed a robust reduction of the 1D5 signal in *TTLL5* mutant ovaries, revealing that *TTLL5* is essential for the appearance of polyglutamylated microtubules in the ovaries and their concentration at the oocyte cortex in S10B.

**Fig. 4.**
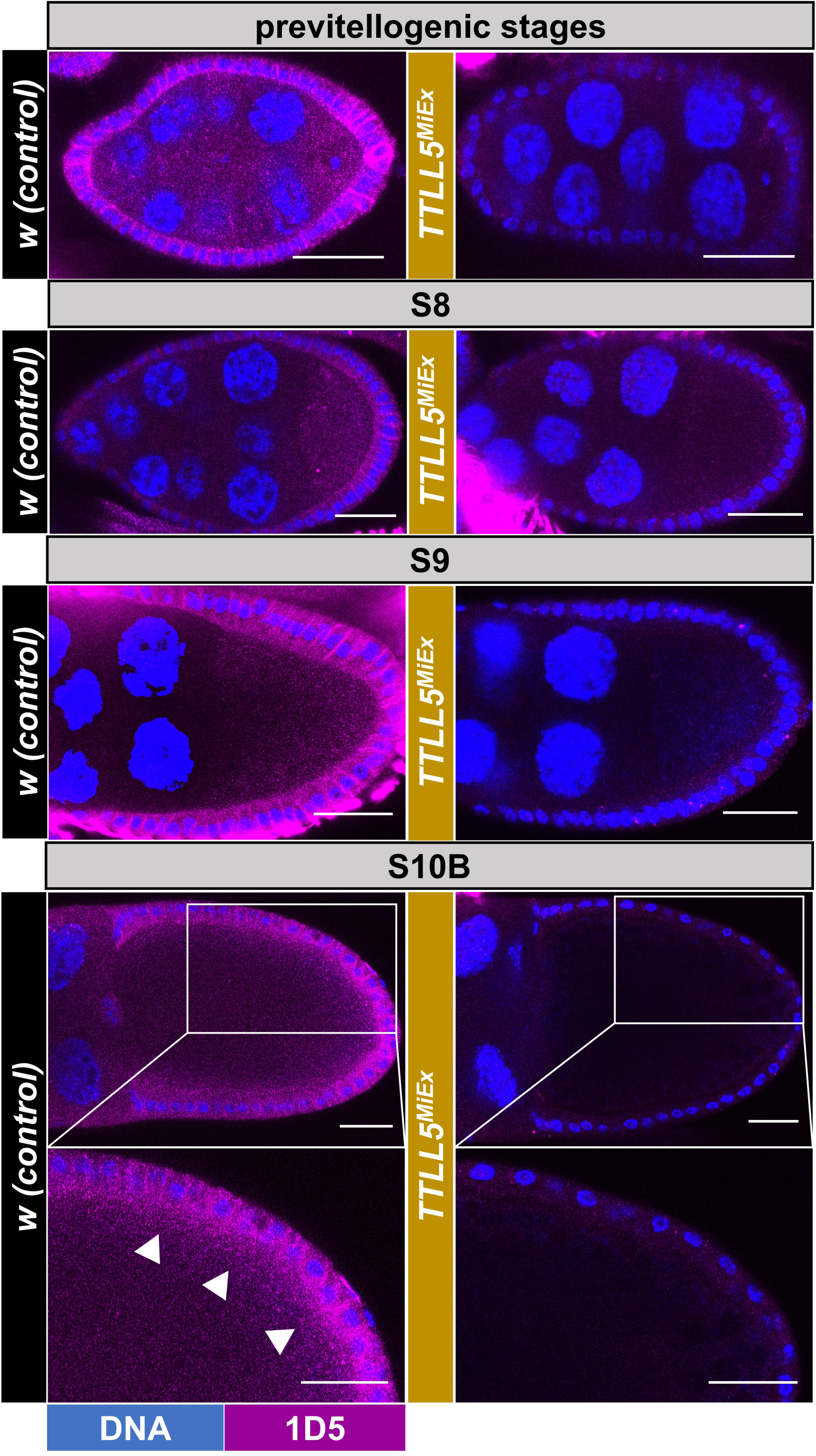
Spatial distribution of polyglutamylated microtubules in the oocyte. Confocal micrographs showing representative oocytes and follicle cells from early to late stages of *w* controls and *TTLL5^MiEx^* mutants. Polyglutamylation (1D5) signals are shown in magenta and Hoechest (blue) stains the DNA. A 1D5 signal is seen in follicle cells and oocytes from previtellogenic to late-stage oocytes. For the S10B oocytes, a high-power magnification of the posterior half is shown, too. At this stage, a slightly biased 1D5 signal intensity was seen in the cortical region of the oocyte (marked by white arrowheads). The 1D5 signal is virtually absent in the *TTLL5* mutants. Imaging conditions and confocal microscope settings were identical for the two genotypes. The genotype of the control was *w.* Genotypes for *TTLL5^MiEx^* were *TTLL5^MiEx^/Df(3R)BSC679*.

### Effect of *TTLL5* on the refinement of posterior localization of Staufen

*osk* mRNA localization, combined with translation control, is essential for targeting Osk protein expression to its proper posterior compartment in the cell during mid and late oogenesis (Weil, 2014). Staufen is a protein that associates with *osk* mRNA into a ribonucleoprotein (RNP) complex during *osk* mRNA localization. It therefore also serves as a proxy for *osk* mRNA localization. Staufen/*osk* mRNA localization requires the dynein/dynactin-mediated minus-end transport machinery in the early stages of oogenesis. Subsequently, the Staufen/*osk* mRNA becomes localized to the posterior of the oocyte from the mid to late stages of oogenesis. This particular Staufen/*osk* mRNA localization phase within the oocyte depends on kinesin-driven processes including active transport and ooplasmic streaming in S10B oocytes (Kato and Nakamura, 2012; Lu et al., 2018; Brendza et al., 2000; Zimyanin et al., 2008).

To determine whether *TTLL5* contributes to these processes, we studied the role of *TTLL5* in the localization of Staufen in ovaries lacking functional TTLL5. In the wild type, the localization of Staufen appeared very tight on the cortex with little lateral extension in S10B (**Fig. 5A**). In ovaries lacking functional TTLL5, we observed a more diffuse localization pattern of Staufen in the S10B egg chambers of all *TTLL5* null mutants (**Fig. 5B**). Z-stack images revealed a more diffusely spread Staufen signal in the *TTLL5* null mutants, although this varied more between different samples. The increased length of the crescent was fully reversed to the regular length in all UAS-*Venus::TTLL5* rescued strains **(Fig. 5C)**. These results suggest that *TTLL5* is required for tight cortical accumulation of Staufen in S10 oocytes. We also analyzed the distribution of Venus::TTLL5 during oogenesis. Antibody staining suggested that Venus::TTLL5 strongly accumulated in the oocyte, especially along the cortex **(Fig. 5C)**. To exclude possible artifacts of the antibody staining, we also tried to observe the self-fluorescent of the Venus tag. Though the Venus fluorescence was relatively weak, Venus::TTLL5 also showed a cortical preference in S10B oocytes, similar to that found by antibody staining (**Fig. S3**). This stronger cortical accumulation of Venus::TTLL5 coincides also with the region where polyglutamylated microtubules were enriched at the same stage **(Fig. 4)**.

**Fig. 5.**
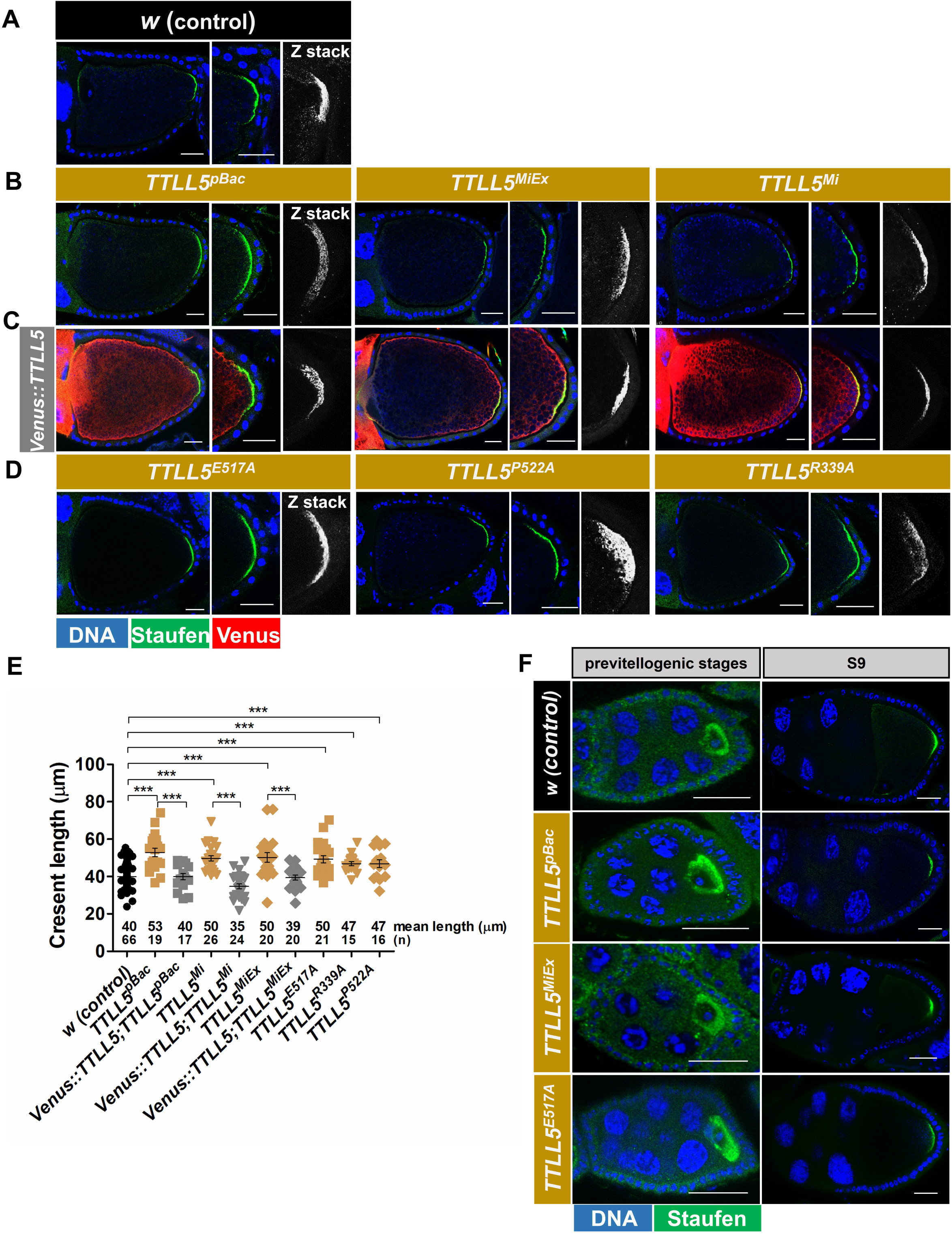
Effect of *TTLL5* on Staufen localization refinement in oocytes. A)-D) Confocal micrographs showing S10B oocytes of *w* controls (A), *TTLL5^pBac^, TTLL5^Mi^*, and *TTLL5^MiEx^* mutants (B), *TTLL5^pBac^, TTLL5^Mi^*, and *TTLL5^MiEx^* mutants rescued by *Mattub4>UASP-Venus::TTLL5* (C), *TTLL5^E517A^, TTLL5^P522A^,* and *TTLL5^R339A^* mutants (D). Anti-Staufen is shown in green, anti-GFP in red, and Hoechst (DNA) in blue. E) Quantification of the Staufen crescent length along the posterior cortex based on A-D) *** p<0.005. z-stack: 12μm. F) Confocal micrographs showing previtellogenic and S9 egg chambers of *w* controls, *TTLL5^pBac^, TTLL5^MiEx^,* and *TTLL5^E517A^* mutants. Staufen signal is shown in green, and Hoechst in blue. Scale Bar: 25μm. *TTLL5^MiEx^* and *TTLL5^E517A^* were hemizygous over *Df(3R)BSC679*. The genotype for *TTLL5^pBac^* was *MattubGal4/+; TTLL5^pBac^/Df(3R)BSC679.* The genotypes of the rescued animals were *MattubGal4/UASP-Venus::TTLL5; TTLL5^alleles^/Df(3R)BSC679*.

In the *TTLL5* point mutants, where specifically the enzymatic activity of TTLL5 was blocked (*TTLL5^E517A^*, *TTLL5^P522A^*, *TTLL5^R339A^*) the localization pattern of Staufen was not refined and the Staufen signal remained less tightly focused (**Fig. 5D**). An established method to quantify the localization of Staufen to the posterior cortex is to measure the length of the posterior Staufen crescent along the cortex (Lu et al., 2016). A shorter crescent is a sign of tighter localization. Indeed, we found that the length of the posterior Staufen crescent was significantly increased in *TTLL5* null and *TTLL5* point mutant ovaries compared to the control and UAS-*Venus::TTLL5* rescued ovaries (**Fig. 5E)**.

We did not observe clear differences in Staufen distribution between the wild type and *TTLL5* mutants in S9 and earlier, previtellogenic stages of oogenesis (**Fig. 5F**). Altogether, these results suggest that the requirement for *TTLL5* for tight cortical accumulation of Staufen starts around S10 and the proper refinement of Staufen localization requires glutamylation of α-tubulin.

### The onset of fast ooplasmic streaming requires *TTLL5*

Ooplasmic streaming is a process that contributes to the refinement of the posterior localization of Staufen (Lu et al., 2016). It is a bulk movement of the oocyte cytoplasm that circulates and distributes mRNAs and proteins in the oocyte, including the Staufen/*osk* mRNA RNPs. It allows these mRNAs and proteins to reach their intended position, where they become anchored. Ooplasmic streaming can be observed in the mid-stage to the late-stage oocytes. Slow and nondirectional flows initiate at stages 8–9 and are followed by rapid and circular streaming at S10B (Lu et al., 2016). The onset of the fast streaming phase has so far been reported to be attributed to 1) microtubule sliding between stably cortically anchored microtubules and free cytoplasmic microtubules close to the cortex (Lu et al., 2016) and 2) subcortical dynamic microtubule-mediated cargo transport (Monteith et al., 2016). Because the final posterior localization refinement of Staufen in S10B oocytes depends on fast streaming (Lu et al., 2016) and also *TTLL5*, we thus wanted to determine if *TTLL5* contributes to fast ooplasmic streaming.

We followed the streaming flow in real-time by measuring vesicle movement in DIC time-lapse movies (**Mov. S1-S5**). We then tracked the movement of the vesicles along the posterior cortical region where microtubule sliding and transport occur (**Fig. 6A**). The kymographs of the tracked vesicles, their analysis, quantification, and interpretation, are shown in **Figs. 6B-6D**. All control ovaries had a typical streaming pattern, and the vesicles were moving unidirectionally parallel to the cortex with a streaming center in the central part of the oocyte. However, 76% *TTLL5^MiEx^* and all *TTLL5^E517A^* mutant ovaries displayed abnormal streaming patterns. These abnormal streaming patterns included a partial disruption of streaming especially in the posterior part of the oocyte and a disordered streaming flow in the entire oocyte. The typical streaming pattern was restored in 71% *TTLL5^MiEx^* ovaries expressing a *Venus::TTLL5* rescue construct. We also evaluated streaming flows in oocytes overexpressing *Venus::TTLL5.* Elevated TTLL5 levels seemed to affect the streaming, too. 29% of the oocytes displayed an abnormal streaming movement or a weak streaming flow like the one observed in *TTLL5* mutants. Moreover, even in oocytes with a circular streaming pattern, the streaming center frequently appeared at a more anterior position in the oocyte, while the cytoplasmic flow in the posterior region was less clearly directed (see **Fig. 6B #2** and **Movie S5 Sample 2**). We thus conclude that the onset of unidirectional fast streaming in S10B oocytes requires moderate levels of *TTLL5*. Again, because abnormal streaming was observed when the enzymatic glutamylation activity of TTLL5 was impaired (*TTLL5^E517A^*), our results suggest that TTLL5 acts on the ooplasmic streaming through its role in MT glutamylation.

**Fig. 6.**
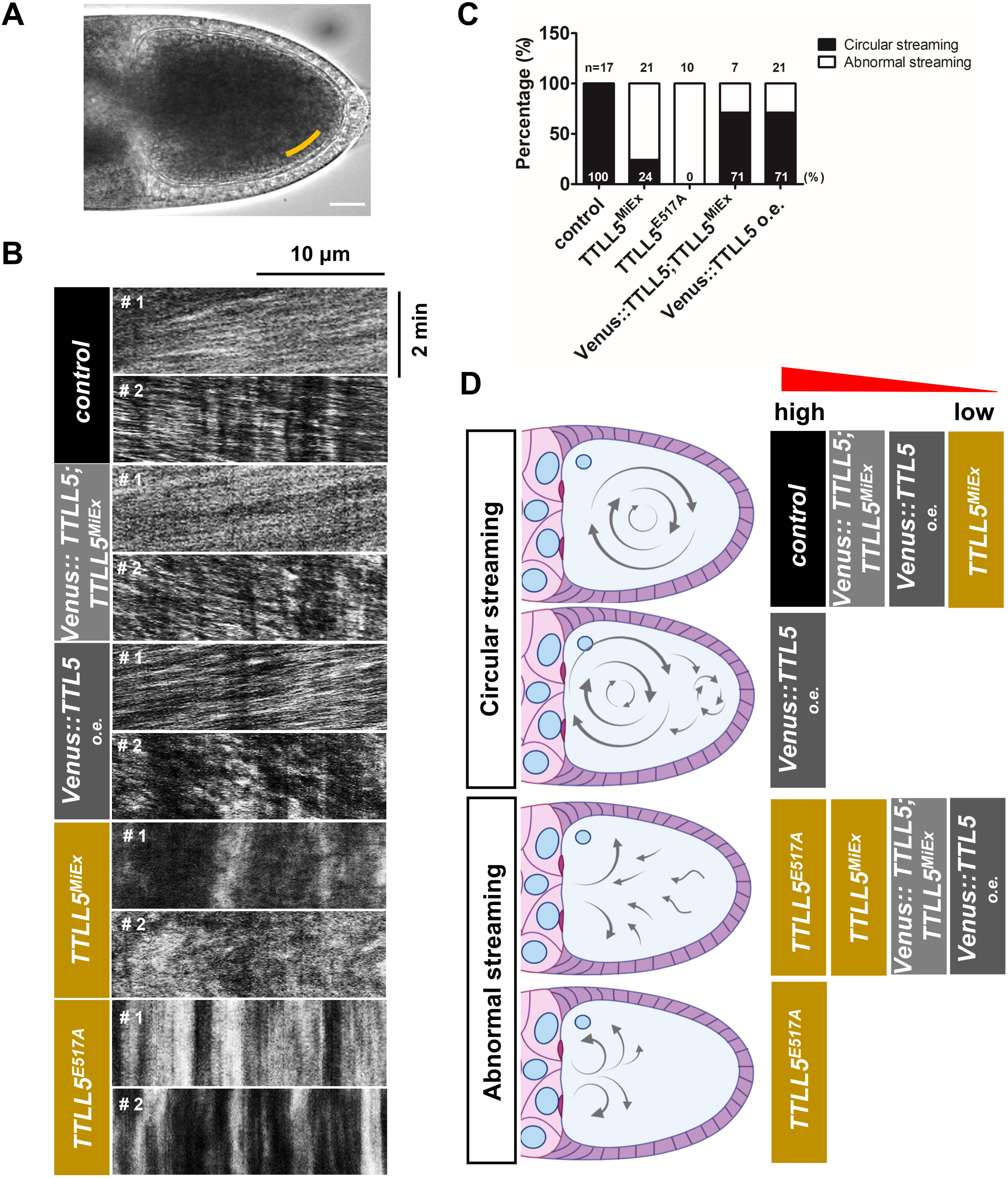
Role of *TTLL5* in fast ooplasmic streaming. A) Example of an S10B oocyte used for particle flow measurements based on time-lapse movies of ooplasmic streaming (**Mov.S1-S5**). Scale Bar: 25 μm. B) Kymographs were generated along a line close to the posterior cortex on one side of the ooplasm (region marked by a yellow line in A). Sample kymographs were shown based on the **Movie S1-S5**. Streaming was shown from 2 representative oocytes for each movie. The first and second oocytes are marked as sample 1 (#1) and sample 2 (#2), respectively. The unidirectional streaming along the posterior cortex is seen in the two control and the *TTLL5* rescue samples, and in sample 1 of the *TTLL5* overexpressing oocyte. Disordered streaming is seen in the second sample of the *TTLL5* overexpressing line and both samples of the *TTLL5* mutants. C) Quantification of streaming patterns seen in the different genotypes. 100% control (n=17), 71%*TTLL5* rescued (n=7) and 71% *TTLL5* overexpression (n=21) S10B oocytes showed an overall circular unidirectional streaming pattern. In contrast, 76% *TTLL5^MiEx^* (n=21) and 100% *TTLL5^E517A^* (n=10) oocytes showed an abnormal streaming pattern. The stronger penetrance of the *TTLL5^E517A^* phenotype (C) compared to the null mutant could be seen as evidence for a slight dominant effect of the point mutation. This is consistent with the stronger expressivity of the streaming defects observed in the movies (B, D), although the numbers appear a bit small to firmly conclude this. D) Schematic representation of the different streaming patterns observed. The frequency with which each streaming pattern was observed in the different genotypes is indicated from high (left) to low (right). “Circular streaming pattern” was further subdivided into “central streaming pattern (top)” and “anterior-biased streaming pattern (below)”. The “central streaming” was observed in all controls, 71% of the *TTLL5* rescue, 43% of the *TTLL5* overexpression, and 21% of the *TTLL5^MiEx^* oocytes. The “anterior-biased streaming” was frequently seen in 28% of the *TTLL5* overexpression ovaries, where the main circular center moved to the anterior part, leaving the posterior with a chaotic streaming flow. “Abnormal streaming” included the oocytes that showed an overall chaotic flow direction (upper one) and the partially disrupted flow (below). The abnormal streaming patterns were mainly seen in the situations when *TTLL5* was insufficient or inactive, and at a low frequency also in *TTLL5* rescued and *TTLL5* overexpressing ovaries. The genotype of the control was *w* or *+/Df(3R)BSC679.* The genotypes for *TTLL5^MiEx^* and *TTLL5^E517A^* were both over *Df(3R)BSC679*. The genotype of the rescued flies was *MattubGal4/UAS-Venus::TTLL5; TTLL5^MiEx^/Df(3R)BSC679.* The genotype of *TTLL5* overexpressing flies was *MattubGal4/UAS-Venus::TTLL5*.

### *TTLL5* affects kinesin distribution in the late-stage female germline

Both Staufen/*osk* mRNA localization and ooplasmic streaming depend on kinesin-1-driven processes (Lu et al., 2016) (Serbus et al., 2005) (Brendza et al., 2000). Therefore, we examined whether *TTLL5* somehow affects Kinesin-1. Firstly, the expression levels of ovarian Kinesin heavy chain (Khc) were neither affected by lack of *TTLL5* function (**Fig. 7A**) nor by *TTLL5* overexpression (**Fig. 7B**). Next, we determined the localization of Khc in stage 10B, when the fast streaming starts. Even though the MT arrays start to disassemble and re-organize in the S10B oocyte, Khc showed a biased enrichment at the posterior in most wild-type control oocytes (**Fig. 7C**). Out of 37 wild-type oocytes, 76% showed a posteriorly enriched Khc signal (**Fig. 7D**). In contrast, Khc distribution showed a posterior oocyte enrichment in only 13% and 14%, respectively, in *TTLL5^pBac^* and *TTLL5^MiEx^* mutant egg chambers, respectively (**Fig. 7C and 7D**). Elevated expression levels of *TTLL5* seemed to affect Khc posterior accumulation as well because a smaller fraction of oocytes (54%) displayed clear posterior Khc enrichment than in the control (**Fig. 7C and 7D**). These results demonstrate that moderate levels of *TTLL5* are needed for the proper posterior enrichment of Khc in S10B oocytes.

**Fig. 7.**
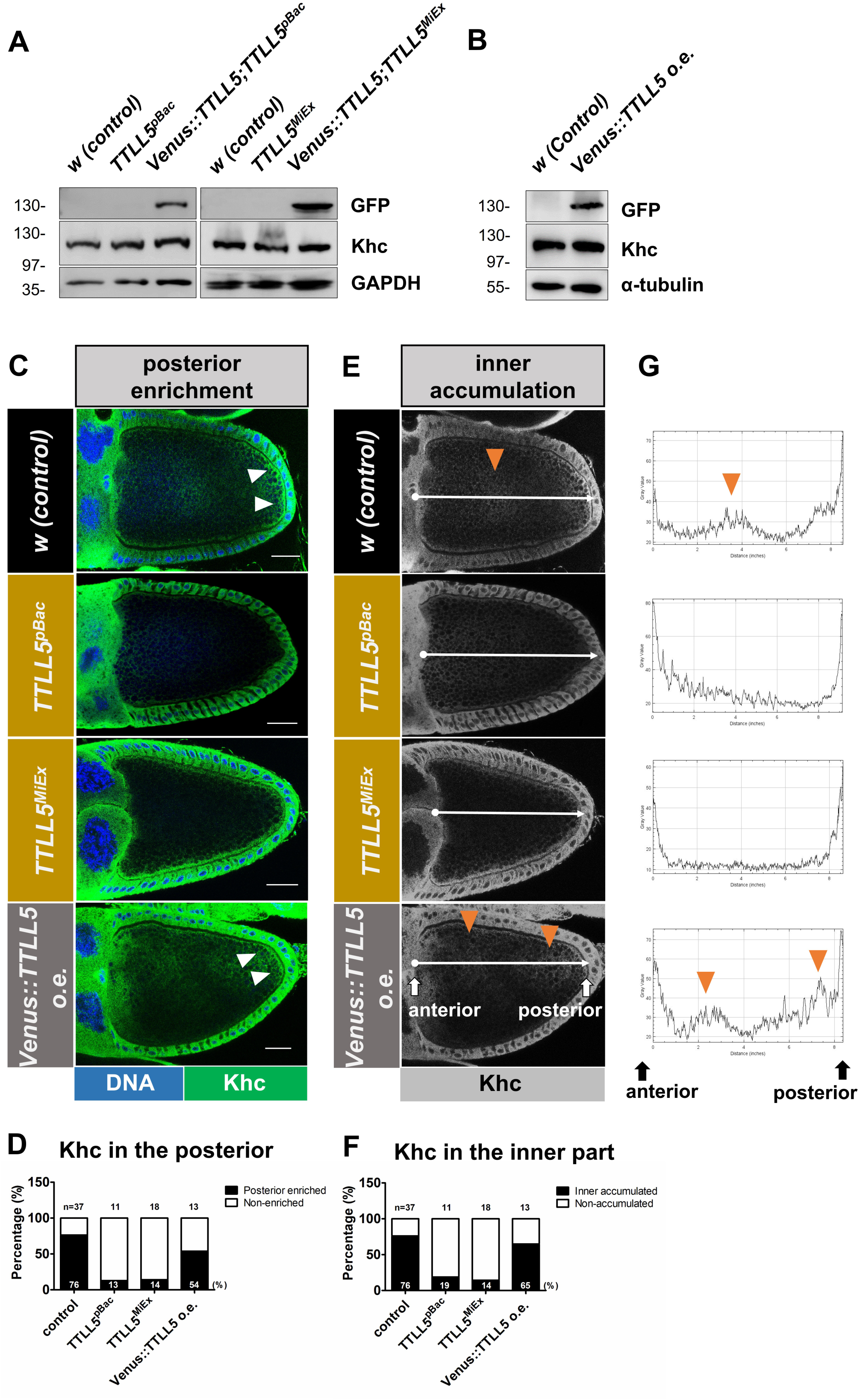
Role of *TTLL5* for kinesin distribution in ovaries. A), B) Khc levels remained unchanged between *w* control, *TTLL5* deficient mutants, rescued *TTLL5* mutants, and *TTLL5* overexpressing ovaries. Anti-GFP antibodies reveal the expression of *Venus:TTLL5.* GAPDH served as the loading control. C) E) Confocal micrographs showing the Khc distribution in stage 10B oocytes. C) White arrowheads point to the posterior enrichment of Khc along the cortical region in the wild-type control and TTLL5 overexpressing oocytes. D) Blind quantification of the posterior enrichment of Khc. The frequency of oocytes with posterior enrichment is shown graphically and numerically. n= number of oocytes evaluated. E) A white line was drawn along the anterior-posterior axis of the oocyte using Fiji. The lines start at the nurse cell oocyte border on the left and end at the posterior follicle cells. The line width was 50 units. G) Intensity charts were plotted based on the line drawn along the AP axis in E). The orange arrowheads in the oocytes and the intensity charts point to the regions showing higher Khc accumulation in the inner region of the oocyte. F) Fraction of the oocytes that showed inner hyperaccumulation of Khc were quantified. n= number of oocytes evaluated. The percentage of oocytes showing inner enrichment of Khc is shown graphically and numerically. Judging Khc distribution was done blindly by two persons. The fractions were taken from the mean values obtained from the two persons. Scale Bar: 25μm. The Khc signal is shown in green, and Hoechst in blue. The genotype for controls was *w* or *+/Df(3R)BSC679.* Genotype for *TTLL5^pBac^*: *MattubGal4/+, TTLL5^pBac^/Df(3R)BSC679*; *TTLL5^MiEx^: TTLL5^MiEx^/Df(3R)BSC679*. Genotype for *TTLL5* overexpression: *MattubGal4/UAS-Venus::TTLL5*.

Khc showed an additional higher accumulation in the inner part of the oocytes (arrowhead in **Figs. 7E**) and this was seen in 76% of the oocytes (**Fig. 7F**). We measured the fluorescence intensity of the Khc signal along the anterior-posterior (AP) axis of the oocytes (line in **Fig. 7E)**. The signal intensity showed a peak in the center or more posteriorly in wild-type oocytes (**Fig. 7G and S4**), indicating that Khc concentration is higher and Khc might function in these regions of the oocyte. In the *TTLL5* mutants, the inner accumulation of Khc was strongly reduced or absent (**Fig. 7E, 7G, S4**). Only 19% of *TTLL5^pBac^* and 14% of *TTLL5^MiEx^* oocytes showed a slight inner Khc accumulation (**Fig. 7F**). Overexpressing TTLL5 in wild-type oocytes showed an inner peak of Khc accumulation in 65%, which is slightly less than the wild-type control. Besides, the center of the single peak appeared in a more anterior position (**Fig. S4**). Interestingly, the higher levels of *TTLL5* also led to the appearance of two peaks of internal Khc accumulation in the oocyte, an anterior and a posterior one (**Fig. 7E, 7G, and S4**). These peculiar inner accumulation patterns of Khc upon *TTLL5* overexpression might explain the unusual ooplasmic streaming patterns with a more anterior center and frequent additional, more posterior streaming center, that appeared in oocytes upon *TTLL5* overexpression. Consistent with the requirement of maintaining the normal circular fast streaming, the proper Khc distribution also needed moderate levels of *TTLL5*. This suggests that Khc is causally connected to the fast cytoplasmic streaming, and both Khc localization and Khc-mediated streaming need *TTLL5*.

We also evaluated the localization of the Bicaudal-D (BicD) protein during the early stages of oogenesis. BicD is the linker that couples diverse cargos to the dynein/dynactin motor and can thus serve to evaluate the dynein transport (Claussen and Suter, 2005). The expression levels of BicD were not affected by the TTLL5 levels (**Fig. S5A**). Also, BicD localized to the posterior of the oocytes, and the ratio of the oocytes showing posteriorly localized BicD was similar in controls and *TTLL5* mutants (**Fig. S5B and S5C**). In conclusion and as suggested by the normal appearance of the egg chambers (see next paragraph), at least the dynein/dynactin/BicD transport on early oogenesis microtubules does not appear to depend on *TTLL5*.

### Female fertility and ovarian development do not require *TTLL5*

The dramatic loss of polyglutamylated ovarian α-tubulin and the phenotypes during late oogenesis in *TTLL5* mutants incited us to analyze whether the morphology of the egg chambers from different stages and female fertility were affected in *TTLL5* mutants. However, the lack of *TTLL5* neither caused a significant decrease in hatching rates, nor distinguishable morphological changes in ovaries. The overall MT polarity of stage 10B oocytes was normal in *TTLL5* mutants based on the localization of the polarity markers Gurken and Staufen protein (**Fig. S6**) (Neuman-Silberberg and Schüpbach, 1996; Zimyanin et al., 2007).

### The glutamylation activity of *TTLL5* modulates the pausing of anterograde axonal transport of mitochondria

Previous studies in *Drosophila* reported that the absence of α-tubulin glutamylation did not seem to be detrimental to the nervous system functions tested in *Drosophila* (Devambez et al., 2017). The wing nerve is an additional, attractive system to monitor organelle transport, such as the movement of mitochondria (Vagnoni and Bullock, 2016) (Hollenbeck and Saxton, 2005). We thus used wing nerves to study the role of *TTLL5* and glutamylation for neuronal transport in the peripheral nervous system by tracking the movement of GFP-labelled mitochondria (Mito::GFP) in the arch of the L1 region which contains bundles of axons (Vagnoni and Bullock, 2016) (**Fig. 8A**). The mitochondria transported towards the thorax (to the right) move by plus-end transport, which is driven by kinesin, whereas mitochondria transported oppositely are transported by dynein (Vagnoni and Bullock, 2016). Time-lapse videos of Mito::GFP movements are shown in **Mov. S6**. Mitochondria are usually transported unidirectionally over a 3 min tracing time. Bidirectionally moving particles were seen, but only rarely. In these cases, they usually moved a short distance toward one direction, suddenly turned back, and ran in the opposite direction. A schematic running pattern is shown in **Fig. 8B**. The start of a run was chosen at the time point when a motile particle began to move away from a stationary position or when it entered the focal plane, and the stop of a run was set when the particle terminated moving or moved out of the focal plane in the given time window. The “pausing” was called when the particle moved with less than 0.2 μm/s within the time window of a “run”. Particles with speeds higher than 0.2 μm/s were considered “trafficking”. Motile mitochondria were manually tracked in both anterograde and retrograde directions. For the anterograde transport, the run length of mitochondria was similar for each “run” between wild-type controls and *TTLL5^E517A^* (**Fig. 8C**). However, a significant increase was observed in the net velocity in *TTLL5 ^E517A^* compared to wild-type controls (**Fig. 8D**). We investigated the reason and found that the absence of enzymatically active *TTLL5* did not alter the trafficking velocity of mitochondria (**Fig. 8E**), but dramatically decreased the total pausing time (1.4 times less) of the anterograde transport of mitochondria (**Fig. 8F**). This suggested that *TTLL5* might not directly affect the trafficking but rather the pausing of mitochondrial transport. To further study the pausing behavior of mitochondria, kymographs of individual motile mitochondria were generated. Kymographs of three representative anterograde transported mitochondria are shown for both genotypes (**Fig. 8G**). The pausing of the mitochondria is pointed out by yellow arrowheads. More frequent pausing was observed in controls compared to the *TTLL5^E517A^*. By analyzing the pausing number in the given distance, we identified a 1.5 times more frequent pausing in controls compared to the *TTLL5^E517A^* mutant (**Fig. 8H**). These results indicate that the longer total pausing time in controls was most likely due to more frequent pausing during anterograde transport. The effect of the lack of TTLL5 enzymatic activity on retrograde transport was less clear because we did not observe significant differences measuring the parameters of the retrograde transport. To sum up, these results suggest that the glutamylation activity of TTLL5 is needed for controlling the pausing of this anterograde axonal transport.

**Fig. 8.**
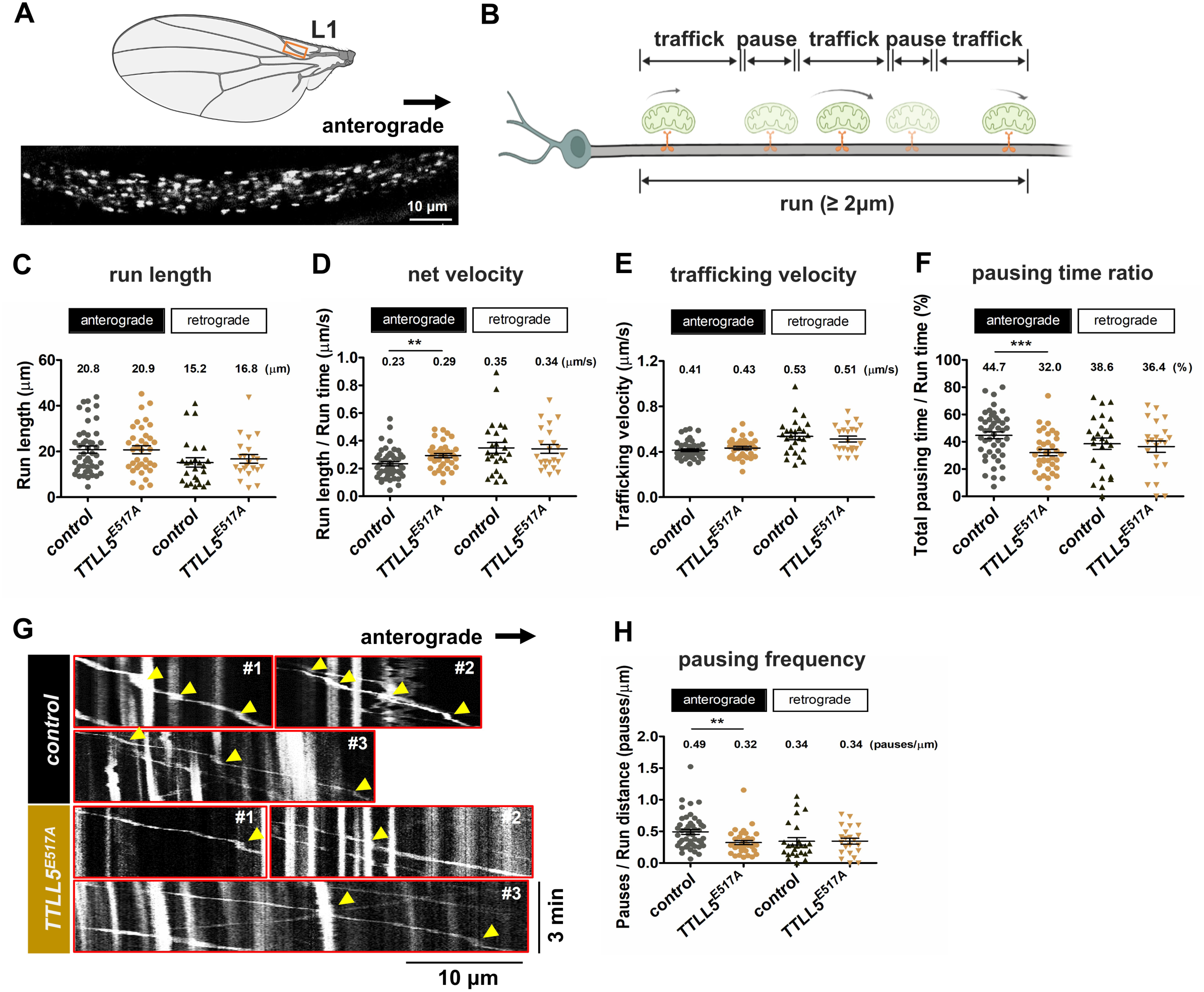
Role of *TTLL5* in the transport of mitochondria in the L1 wing nerve. A) Representative still image of GFP-labelled mitochondria in axons of the L1 vein of both control and *TTLL5^E517A^* wings. The position of the L1 region is indicated in the picture above. B) Scheme of the transport behavior of the mitochondria in the wing neuron. See the main text for a detailed description. The schema was drawn by biorender.com. C-F) Mean values of run length, net velocity, trafficking velocity, and pausing time ratio in the control and the *TTLL5 ^E517A^* mutant for both anterograde and retrograde transport. C) The run length is the sum of the distances traveled by individual mitochondria in a trafficking state. D) The net velocity indicates the run distance divided by the run time. The run time is the sum of the total pausing time plus the total trafficking time. E) Trafficking velocity indicates the mean of the instant velocity of the trafficking mitochondria. F) The pausing time ratio was calculated as total pausing time divided by run time. The total pausing time was the sum of the time when the mitochondria were pausing. G) Kymographs were generated from 3 representative mitochondria in the control and the *TTLL5^E517A^* mutant. Yellow arrowheads indicate the pauses of selected mitochondria over the 3 min time window. H) The mean values of pausing frequency. Pausing frequency indicates the number of pauses divided by run length. Mitochondria were quantified from 7 control fly wings and 8 *TTLL5^E517A^* fly wings. The numbers of mitochondria (n) analyzed for the anterograde control, anterograde *TTLL5^E517A^*, retrograde control, and retrograde *TTLL5^E517A^* were 46, 36, 25, and 21, respectively. *** p<0.005, ** p<0.01. Genotypes for controls were a mixture of *ApplGal4;UAS-Mito::GFP;+/Df(3R)BSC679* and *ApplGal4;UAS-Mito::GFP;TTLL5 ^E517A^/+*. Genotypes for *TTLL5^E517A^* mutants were *ApplGal4; UAS-Mito::GFP; TTLL5^E517A^/Df(3R)BSC679*.

## Discussion

### Glutamylation of the C-terminal tail of ovarian α-tubulin

Side chain glutamylation of the C-terminal domain of α-tubulin in *Drosophila* ovaries was observed using specific antibodies and MS identification. Results from wild-type ovaries and mutants revealed that *TTLL5* is essential for this modification. Consistent with results from mammalian cells and the *Drosophil*a nervous system (Devambez et al., 2017; van Dijk et al., 2007), the strongly reduced or absent Western blot signal of monoglutamylated tubulin in the presumptive null mutants of *TTLL5* **(Fig. 3B)** shows a requirement for *TTLL5* for the accumulation of α-tubulin with the first branching Glu side chain added to one of the C-terminal Glus. The MS analysis led to the same conclusion because it did not uncover any glutamylation in the *TTLL5* mutants. In theory, this could mean that *TTLL5* is required for the addition of this first Glu side chain residue or it might only be required for the stability of glutamylated α-tubulin. TTLL5 has the enzymatic activity to add glutamyl groups and mammalian TTLL5 is considered to be an initiator glutamylase. We can, therefore, assume that the former is the case in *Drosophila*, too. TTLL1 and TTLL6 are elongators of polyglutamylation of α-tubulin (Gadadhar et al., 2017; van Dijk et al, 2007; Janke et al., 2005). However, there is no evidence that the homologous *TTLL6A*/*TTLL6*B or *TTLL1A/1B* are expressed in *Drosophila* ovaries **(Table S1)**. The robust reduction of polyglutamylated tubulin in *TTLL5* mutants observed in ovaries (this work) and the nervous system (Devambez et al., 2017) further shows that *TTLL5* is also needed, directly or indirectly, for polyglutamylation in *Drosophila.* However, we cannot easily figure out *in vivo* if TTLL5 itself has the activity to extend the oligo-Glu side chains. TTLL4A is another enzyme that is expressed in ovaries. Though the predicted mouse homolog of the *Drosophila* TTLL4A prefers to glutamylate β-tubulin (van Dijk et al., 2007; Mahalingan et al., 2020), our data revealed a strong requirement for *Drosophila TTLL4A* for the proper glutamylation of α-tubulin **(Fig. S2C)**, and the Western blotting experiments suggested that it might be required for both monoglutamylation and polyglutamylation **(Figs. S2B and S2C)**. However, the interpretation of the Western blotting results with the GT335 antibody is often difficult and further mass spectrometry analyses should be performed to verify the status of glutamylation of MTs in the mutants.

The MS results revealed that glutamylation strongly preferred the substrate αTub84B/D over αTub67C, even though αTub67C is an ovarian-specific α-tubulin. This points to a possible need for a side-chain modification-resistant α-tubulin in the germline to regulate different transport processes by providing local MT sites with reduced side-chain glutamylation, possibly to allow for longer processive movement of kinesin motors as was observed in the absence of glutamylation in the mitochondrial transport assay **(Fig. 8)**. It will also be interesting to find out whether MT tracks containing specific hypoglutamylated regions exist in oocytes and whether these perform an important function in large insect oocytes that use rapid cytoplasmic streaming to localize patterning factors.

### Function of glutamylation of microtubules on Kinesin-1 dependent streaming in late-stage oocytes

The unidirectional flow seen during rapid ooplasmic streaming is caused by the Kinesin-1-dependent MT sliding and MT cargo transport along the cortical and subcortical MTs (Lu et al., 2016) (Monteith et al., 2016). The proper localization of Khc to the posterior cortical region **(Fig. 7)** and the glutamylation of cortical MTs **(Fig. 4**) depend on TTLL5, which is normally also concentrated at the cortex in S10B oocytes (**Fig. 5C** and **Fig. S3**). The observation of these correlations in the cortical region point to the glutamylation of the MTs as a precondition for the proper localization and function of Kinesin-1.

Kinesin-1 also accumulates at higher levels in the inner part of the oocyte (**Fig. 7**), suggesting that the inner-oocyte MTs might also contribute to the rapid streaming observed. This hypothesis is supported by the observation that hyperglutamylation by TTLL5 overexpression often leads to an anterior-biased accumulation of Khc in the interior of the oocyte or the formation of two inner Khc clusters along the anterior-posterior axis (**Figs. 7E, G, Fig. S4**). This abnormal Khc distribution is then mirrored by the streaming pattern in such oocytes (**Fig. 6D**), suggesting that it may cause the shift of the streaming center to the anterior part of the oocytes and the appearance of a second posterior streaming center. Further investigations through direct tracking of Khc movement in S10B oocytes under hypo- and hyperglutamylation conditions may provide more insight into the impact of MT glutamylation on Kinesin-1 function *in vivo*.

### Role of glutamylation of α-tubulin for stopping transported mitochondria

The previous study showed that α-tubulin was the only tubulin subunit that can be glutamylated in the *Drosophila* nervous system and *TTLL5* is required for α-tubulin glutamylation in the neurons (Devambez et al., 2017). However, the absence of *TTLL5* did not reveal clear defects in larval neuromuscular junction morphology, larval locomotion behavior, or vesicular axonal transport in larval neurons. We, therefore, used a different system, the transport of mitochondria in the adult wing axon to test for a possible neuronal function.

By measuring the parameters related to the pausing time and pause frequency, we found that the normal stopping or pausing of mitochondria during axonal transport needs *TTLL5* and its glutamylase activity, suggesting that glutamylated α-tubulins are needed for this effect. Glutamylated side chains of α-tubulin might act as marks or even obstacles that disturb the movement of mitochondria. Consistent with our results, a previous *Drosophila TTLL5* study mentioned that a slight effect on pausing of vesicles during axonal transport had also been noted in *TTLL5* mutants but turned out to be not significant (Devambez et al., 2017). Additionally, a recent study revealed a similar function of glutamylation of α-tubulin for the transport of mitochondria (Bodakuntla et al., 2021). The absence of polyglutamylation of α-tubulin in *TTLL1^-/-^* increased the motility of mitochondria in the mouse neurons but did not affect their transport velocity. The results led to the hypothesis that glutamylation of α-tubulin may not directly affect transport but rather affect mitochondrial docking and motor binding to MT tracks by acting as roadblocks on the MT tracks.

Our results showed a selective effect of Glu-MTs on mitochondrial motility for plus-end transport in axons, a process that is carried out by Kinesin-1 (Pilling et al., 2006). This suggests that glutamylation of MTs plays a role in controlling Kinesin-1 activity similar to what we had observed in the stage 10B oocyte. Several *in vitro* studies have also shown that Glu-MTs affect Kinesin-1 activity. The considerable negative charge of the modified regions in the poly-Glu-modified tubulins might interfere with the interaction of kinesin with tubulin (**Fig. 1B;** Skiniotis et al., 2004). *In vitro* studies demonstrated that the artificial tethering of glutamyl peptides to the side chains of MTs affected the function of Kinesin-1 (Sirajuddin et al., 2014). More recent *in vitro* studies further reveal that TTLL7-modified β-tubulin polyglutamylation specifically decreases the motility of Kinesin-1 (Genova et al., 2023). All these observations suggest that side chain glutamylation of MTs is sensed by Kinesin-1 and functions to interfere with the MT binding of Kinesin-1, thus disturbing the movement of Kinesin-1 dependent cargos. However, an alternative hypothesis is suggested by the work of Lacroix and colleagues (Lacroix et al, 2010). They showed that glutamylation of MTs leads to the breakage of the “MT road” by recruiting severing proteins and this is then followed by the disruption of cargo transport. These studies showed that the severing rate of a microtubule severing protein Spastin was increased in the presence of MTs that were glutamylated by TTLL4 or TTLL6 in cultured HeLa cells. Another group demonstrated that the glutamylated MTs (modified by TTLL7) increased Spastin-mediated severing when glutamic acid side chains were shorter than 8Es (Valenstein and Roll-Mecak, 2016), which is the case in *Drosophila* ovaries **(Fig. 1)**. A second MT severing protein has recently entered the scene in the same context. The interaction of Katanin with microtubules is also enhanced by α-tubulin tail glutamylation in *C. elegans* (Szczesna et al., 2022). *Drosophila* Spastin is present in the axons and synaptic connections of larvae. However, the effects of its absence in *Drosophila* larval neurons are controversial. One group suggests that the loss of Spastin leads to a reduction in synapse growth and an increase in synaptic transmission (Trotta et al., 2004), while another study reports the opposite effect (Sherwood et al., 2004). *Drosophila* Katanin acts as a severing protein and is required for proper synapse development in the larval neuromuscular junction (NMJ; Díaz-Valencia et al., 2011; Mao et al., 2014). It would also be valuable to investigate the impact of broken MTs on axonal cargo transport using the adult wing axon system, which presents a unique method to study this phenomenon. Because α-tubulin is the major substrate for glutamylation in the *Drosophila* nervous system, this system is a powerful and simple *in vivo* model to study in greater detail the effect of glutamylation of these specific α-tubulin isotypes on the movement of MT motors.

### Physiological relevance of α-tubulin glutamylation

Even though an essential function of α-tubulin glutamylation has not emerged from this study, it has become clear that Kinesin-1-dependent processes are modified or controlled by this modification and that evolution has adapted the tubulin code to optimize the usage of MTs for different evolving needs. That *TTLL5* is not essential for survival also makes this protein an interesting candidate for drugs that can modulate its glutamylation activity to regulate and possibly even increase MT transport in neurodegenerative diseases where MT transport is known to be affected (Bodakuntla et al., 2020)

## Materials and methods

### *Drosophila* genetics

The fly stocks used in this study, *MattubGal4* (7062), *TTLL5^pBac^* (16140; allele name *TTLL5^B093^*), *TTLL5^Mi^* (32800; allele name *TTLL5^MI01917^*), and *Df(3R)BSC679* (26531) were obtained from the Bloomington Drosophila stock center. The *TTLL5^MiEx^* mutation was made by imprecise excision of the Minos element, which left a 3-base insertion (CTA), which introduced a premature stop codon at the genomic position 8,132 in the open reading frame (polypeptide position 392). The different *TTLL5* point mutations were generated by CRISPR/Cas9. The two *Drosophila* stocks used for *TTLL5* gene editing, *y[1] M{w[+mC]=nos- Cas9.P}ZH-2A w[*]* and *y[1] v[1] P{y[+t7.7]=nos-phiC31\int.NLS}; P{y[+t7.7]=CaryP} attP40* were obtained from the Bloomington *Drosophila* Stock Center (54591) (Port et al., 2014) and (25709), respectively. Target sequences near the K131, R339 R376, and E517 codons were selected to introduce INDELs and the following point mutations: K131A, R339A (b6-7 loop), R376A, and E517A. Phosphorylated and annealed primers (**Table S3A**) needed to construct the corresponding coding sequences for a single guide RNA (sgRNA) were each cloned into the BbsI site of *pCFD5* according to the protocol published on www.crisprflydesign.org. After amplifying the vector in XL1 blue cells, the plasmids were sequenced to verify the correct assembly. Subsequently, they were injected into *attP40* embryos. *TTLL5*-guide stocks with integrated genes encoding sgRNAs were established *(=v; attP40{v^+^, TTLL5-guide}/CyO*). To obtain INDELs for TTLL5, a three-generation crossing schema was set up. Firstly, *nos-cas9* virgins were crossed with males from the different TTLL5-guide stocks. Secondly, the subsequent single male progeny was crossed with *w; Ly/TM3, Sb* virgins. Thirdly, resulting in single *TM3, Sb* males were again crossed with *w; Ly/TM3, Sb* virgins to establish the stocks. The nature of the mutated *TTLL5* alleles was determined by sequencing the targeted region in each stock and comparing it to the sequence of the *TTLL5* gene of the parental *y w {w^+^, nos-cas9}* mothers and *TTLL5-guide* fathers (**Table S3B**). All the *TTLL5* alleles were crossed to *Df(3R)BSC679* to analyze the effect of the *TTLL5* mutations in hemizygous animals. Abnormally shaped egg chambers were seen in *TTLL5^pBac^/Df(3R)BSC679.* But it seems to be the second site effect as the phenotype disappeared when it was in combination with specific 2**^nd^** chromosomes, for example, *MattubGal4/+ or +/UAS-Venus::TTLL5)*.

The *UASp-venus::TTLL5* DNA construct was made from the *TTLL5* cDNA amplified by PCR (forward primer: 5′ - GTTCAGATCTATGCCTTCTTCATTGTGTG -3′; reverse primer: 5′ -GAGTCATTCTAGAGCTTCATAGAAATACCTTCTCC- 3′). The resulting DNA was inserted into the *pUASP-venus* vector via Xba l/Bgl II ligation and targeted into the *attP-58A* landing platform (24484) to generate the transgenetic *UASp-venus::TTLL5* fly strain. *ApplGal4* and *UAS-mito::GFP* were kindly provided by Simon Bullock’s group (Vagnoni et al., 2016).

The *TTLL4A* loss-of-function mutants were *TTLL4A^pBac^* (17925; allele name *TTLL4A^e01119^*), *TTLL4A^pEPgy^* (22659; allele name *TTLL4A^EY23195^*), and *Df(2L)BSC145* (9505). They were obtained from the Bloomington *Drosophila* stock center. The *TTLL4A^pBac^* strain carries a piggyBac transposon that had inserted into the leader sequence of *TTLL4A*. The *TTLL4A^pEPgy^* carries a pEPgy transposon that had inserted into the promotor region of *TTLL4A.* The *Df(2L)BSC145* removes the region that covers *TTLL4A*.

### Western blotting

Western blotting performed with ovarian extracts was carried out with 10-20 pairs of ovaries for each genotype. Newly hatched females of the indicated genotypes were collected and crossed with *w* males in fresh vials with ample yeast food for two to three days. Fully developed ovaries were then dissected and collected. The samples were directly lysed in Laemmli buffer, boiled at 95°C for 10 min, fractionated by SDS-PAGE, and transferred onto PVDF membranes. The following primary antibodies were used: mouse 1D5 (1:500, Synaptic System), mouse GT335 (1:200, Adipogen), mouse DM1A (1:1,000, Sigma), mouse tyrosinated α-tubulin (1:1,000, Sigma), rabbit GFP (1:2,000, Sigma), rabbit GAPDH (1:1,000, GeneTex), rabbit *Drosophila* Clathrin light chain (Clc) (1:2,000, Heerssen et al., 2008), mouse BicD (1B11 plus 4C2, 1:5) (Suter and Steward, 1991), rabbit Khc (1:1,000, cytoskeleton), mouse TAP952 (1:500, Merck). For testing the expression of TTLL5, the ovaries were homogenized in ice-cold lysis buffer (150 mM NaCl, 50 mM Tris-HCl (pH 7.5),1 mM MgCl_2_,1 mM EDTA, 0.1% Triton X-100, 0.5 mM DTT, and protease inhibitor cocktail (Roche)) and centrifuged at 13,200g for 20 min. The supernatant with the soluble proteins was collected and loaded for WB. The rabbit anti-Drosophila TTLL5 (1:500) was generated by the GenScript company. The fragment used for antigen induction was: TKLLRKLFNVHGLTEVQGENNNFNLLWTGVHMKLDIVRNLAPYQRVNHFPRSYEMTRKDRLYKNIERMQHLRGMKHFDIVPQTFVLPIESRDLVVAHNKHRGPWIVKPAASSRGRGIFIVNSPDQIPQDEQAVVSKYIVDPLCIDGHKCDLRVYVLVTSFDPLIIYLYEEGIVRLATVKYDRHADNLWNPCMHLCNYSINKYHSDYIRSSDAQDEDVGHKWTLSALLRHLKLQSCDTRQLMLNIEDLIIKAVLACAQSIISACRMFVPNGNNCFELYGFDILIDNALKPWLLEINLSPSMGVDSPLDTKVKSCLMADLLTCVGIPAYS. Secondary antibodies were HRP-linked goat anti-mouse (1:10,000, GE Healthcare) and HRP-linked goat anti-rabbit (1:10,000, GE Healthcare). Antibodies were incubated in 5% non-fat milk PBST (PBS with 0.1% Triton X-100) and washed with PBST. Chemiluminescence detection was performed with the Amersham imager 600 (GE Healthcare).

### Mass Spectrometry

Proteins were extracted from the ovaries of 100 3-day-old, well-fed females of each genotype, crossed to wild-type males, and proteins were boiled and run on SDS-PAGE. The gel was then stained with Coomassie Blue. The gel pieces containing the α-tubulin region were cut, alkylated, and digested by trypsin N (ProtiFi) for 3 hours at 55°C. C-terminal peptides of α-tubulin were analyzed by liquid chromatography (LC)-MS/MS (PROXEON coupled to a QExactive mass spectrometer, Thermo Fisher Scientific). Samples were searched against the UniProtKB *Drosophila* database using Pipeline, PEAKS, Easyprot, and MSfragger as search engines. Peptides were identified by PSM value (peptide spectrum match). To avoid repetition of total PSM quantification, we searched only for one type of PTM in an analysis each time. Total PSM indicates the sum of PSMs of primary C-terminal peptides with either glutamylation or glycylation modifications or non-modified peptides.

### Ooplasmic streaming in living oocytes

Three days old females were kept together with wild-type males under non-crowding conditions at 25°C on food containing dry yeast. The determination of ooplasmic streaming was based on the method of Lu and colleagues (Lu et al., 2016). Ovaries were directly dissected in 10 S halocarbon oil (VWR), and egg chambers were teased apart in the oil drop on 24×50 mm coverslips (VWR). Living Stage 10B oocytes were directly observed and imaged under the 60×1.4 NA oil immersion objective on a Nikon W1 LIPSI spinning disk with a 2x Photometrics Prime BSI CMOS camera. Time-lapse images were acquired every 2s for 2 min under bright-field (BF) controlled by Nikon Elements software. Flow tracks were generated by kymographs using Fiji (Schindelin et al., 2012) plugin Multi Kymograph (HTTPS:// imagej.net/ Multi _Kymograph).

### Immunostaining and confocal microscopy

Ovaries were dissected in 1× PBS and fixed in buffer (200 μl 4% formaldehyde (Sigma), 600 ul Heptane (Merck Millipore), 20 ul DMSO (Sigma) for each group) for 20 min. Ovaries were then blocked in 2% milk powder in PBST (PBS, 0.5% Triton X-100). Samples were incubated with rabbit anti-Staufen (1:2,000) (St Johnston et al., 1991), mouse anti-Gurken (1D12, 1:10) (Queenan et al., 1999), mouse anti-BicD (1B11 plus 4C2, 1:5) (Suter and Steward, 1991), rabbit anti-Khc (1:150, Cytoskeleton), rabbit anti-GFP (1:200, Sigma) and mouse anti-1D5 (1:150, Synaptic System) at 4°C overnight, and incubated with secondary Alexa Fluor goat anti-mouse Cy3 (1:400, Jackson ImmonoResearch) and Alexa Fluor goat anti-rabbit 488 (1:400, Life technologies Thermo Fischer) at room temperature for 3h. DNA and nuclei were stained using Hoechst 33528 (Thermo Fischer Scientific) for 10 min. Mounted samples were imaged with a Leica TCS SP8 confocal microscope equipped with a 63×1.40 oil lens and controlled by the Las X software (Leica). Images of single focal planes and 12 μm Z projection were acquired. The crescent length of Staufen was processed by the Fuji segmented line tool based on a single plane. The data were shown as means ± SEM by Prism 5 (GraphPad software). The statistical significance was determined by an unpaired two-tailed non-parametric t-test (Mann-Whitney test). For the Khc intensity analysis, the intensity of fluorescence in the oocyte was plotted and the intensity charts were generated using Fiji’s “Analyze-Plot profile” function.

### Mitochondrial transport in *Drosophila* wing neurons

Mitochondrial transport in *Drosophila* wing neurons was tracked as described (Vagnoni and Bullock, 2016). In brief, 1-2 days old flies were anesthetized with CO_2_ and immobilized in a custom-built chamber formed of two No.1.5 24×50 mm coverslips (VWR). Imaging of wings was performed with a Nikon W1 LIPSI spinning disk microscope equipped with a 2x Photometrics Prime BSI CMOS camera and a 60×1.4 NA oil immersion lens. Time-lapse video of a single focal plane was acquired with an acquisition rate of 0.5 frame/s for 3 min for Mito::GFP using a 488 nm laser.

Quantification of the movements was performed in Fiji. At least 7 wings of each genotype were tracked for quantification. A region of the wing arch of the L1 vein was selected and straightened with the Straighten plugin (Eva Kocsis, NIH, MD). For each “run”, the total trafficking distance of more than 2 μm was defined as “motile mitochondria”. To exclude the oscillatory movements of mitochondria, a minimal instant velocity between each frame was defined at 0.2 μm/s. Transported mitochondria were manually tracked by the manual tracking plugin (Fabrice Cordelières, Institute Curie, France). Kymographs of individual mitochondria were produced using Fiji (Schindelin et al., 2012) plugin Multi Kymograph (HTTPS:// imagej.net/ Multi _Kymograph). Run length, instant velocity values, and total running, trafficking & pausing times were exported into Excel for analysis. The data are shown as means ± SEM by Prism 5 (GraphPad). The statistical significance was determined with the student’s t-test.

### Sequence alignment

Sequences were aligned by CLC Sequence Viewer 8.0 software (Qiagen). The UniProtKB accession numbers are as follows: *HsTTL*: Q8NG68; *HsTTLL5*: Q6EMB2; *MmTTLL5*: Q8CHB8; *DmTTLL5*: Q9VBX5;

## Supporting information

Movie. S1 Streaming of w (Control)

Movie. S2 Streaming of TTLL5 MiEx mutation

Movie. S3 Streaming of TTLL5 E517A mutation

Movie. S4 Streaming of Venus-TTLL5 rescue on TTLL5 MiEx mutation

Movie. S5 Streaming of Venus-TTLL5

Movie. S6 Mitochondrial trafficking in L1 vein

## Acknowledgments

We thank Simon L. Bullock (Cambridge, UK) for *ApplGal4* and *UAS-mito::GFP* fly lines, Michel Steinmetz (Paul Scherrer Institute, Switzerland), and the Carsten Janke (Institute Curie, France) laboratory members for discussions and suggestions. We also thank the University of Bern (Switzerland) Mass Spectrometry center members Sophie Braga Lagache and Natasha Buchs for performing the MS experiments and Manfred Heller and Anne-Christine Uldry for peptide analysis and discussion of the results.

## Support

This work was financially supported by the project grants 31003A_173188 and 310030_205075 from the Swiss National Science Foundation (SNF; https://www.snf.ch/en) to BS. Equipment support was by an equipment grant from SNF (316030_150824) to BS and by the University of Bern (https://www.unibe.ch). Additional salary and material support were received from the University of Bern (https://www.unibe.ch) to BS. The funders had no role in the study design, data collection, analysis, decision to publish, or manuscript preparation.

The authors declare no competing financial interests.

## Author contributions

Mengjing Bao, Beat Suter, and Maria Paula Vazquez-Pianzola conceived the project and designed the experiments. Mengjing Bao, Beat Suter, Ruth Dörig, and Dirk Beuchle performed the experiments and analyzed the data; Mengjing Bao and Beat Suter wrote the manuscript.

---

**Table S1:**
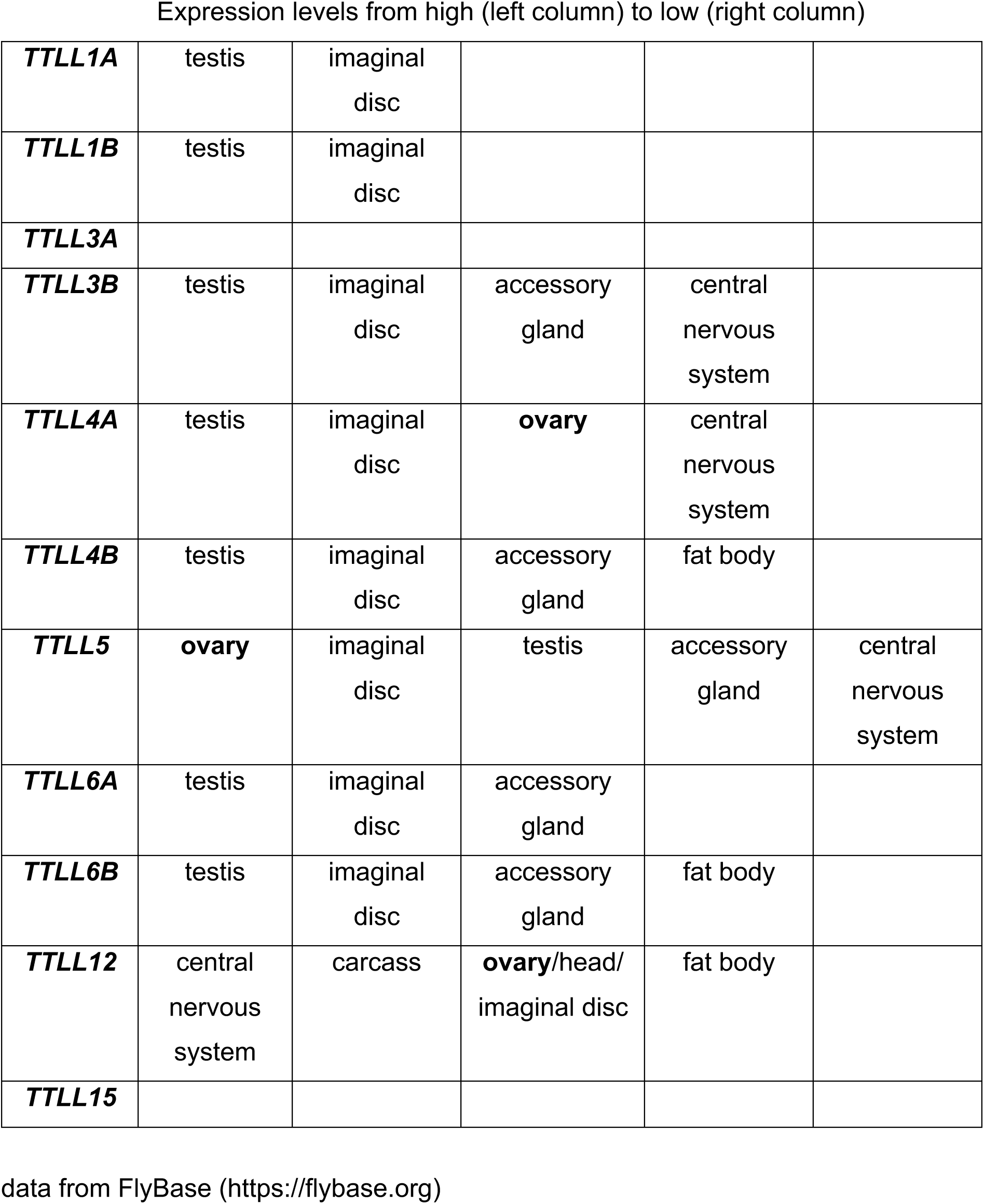
Tissue specific expression of the 11 Drosophila TTLL genes.

**Table S2.** Number of peptides of αTub84B, αTub84D, and αTub67C found by MS.

| **Isoform** | ***w*** | ***TTLL5<sup>pBac</sup>*** | ***TTLL5<sup>MIEX</sup>*** |
| --- | --- | --- | --- |
|  |  | (PSM value<sup>1)</sup>) |  |
| αTub84B/D | 56 | 29 | 7 |
| αTub67C | 12 | 12 | 4 |

<sup>1)</sup> The total peptide numbers were revealed by the values of total PSM (peptide spectrum match) of peptides containing the full primary C-terminal sequence.

**Table S3A:**
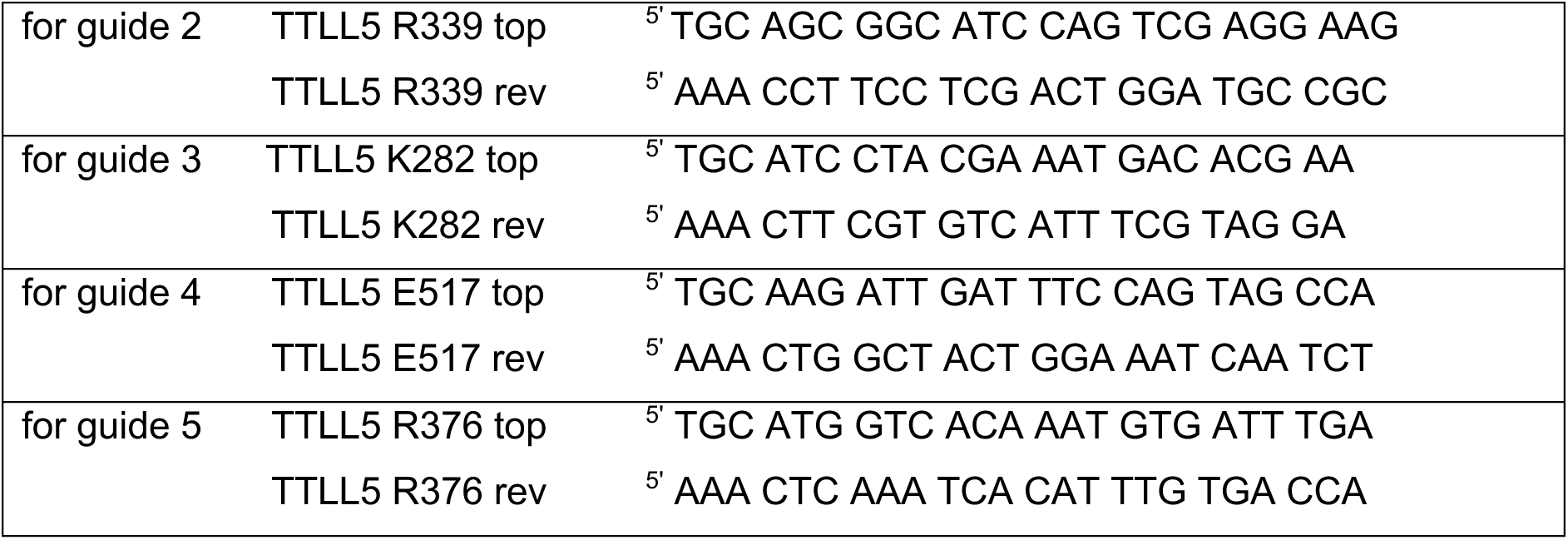
primers for cloning the sequences encoding the sgRNAs into the Drosophila transformation vector pCFD5.

**Table S3B:**
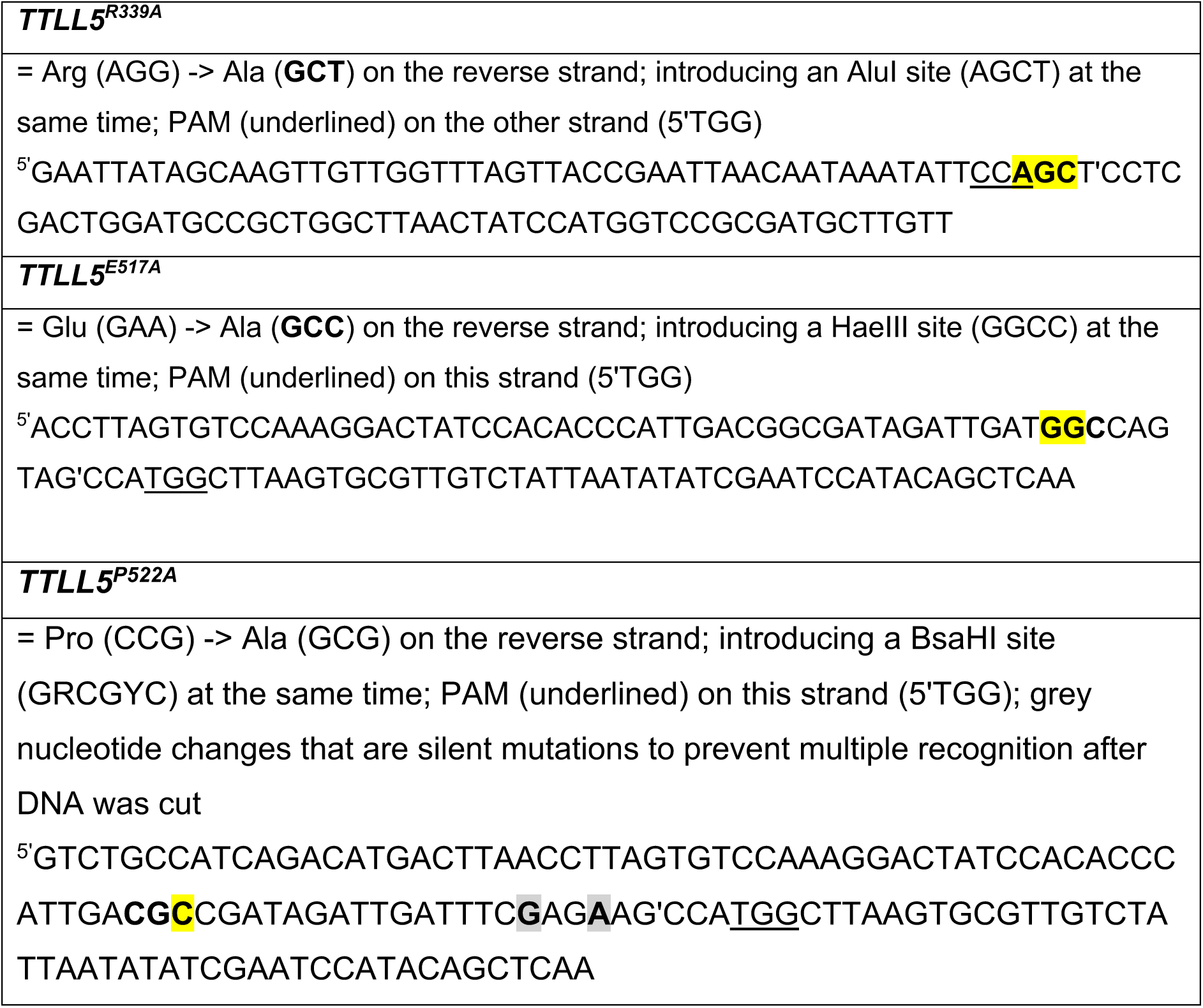
templates to introduce point mutations.

**Fig. S1:**
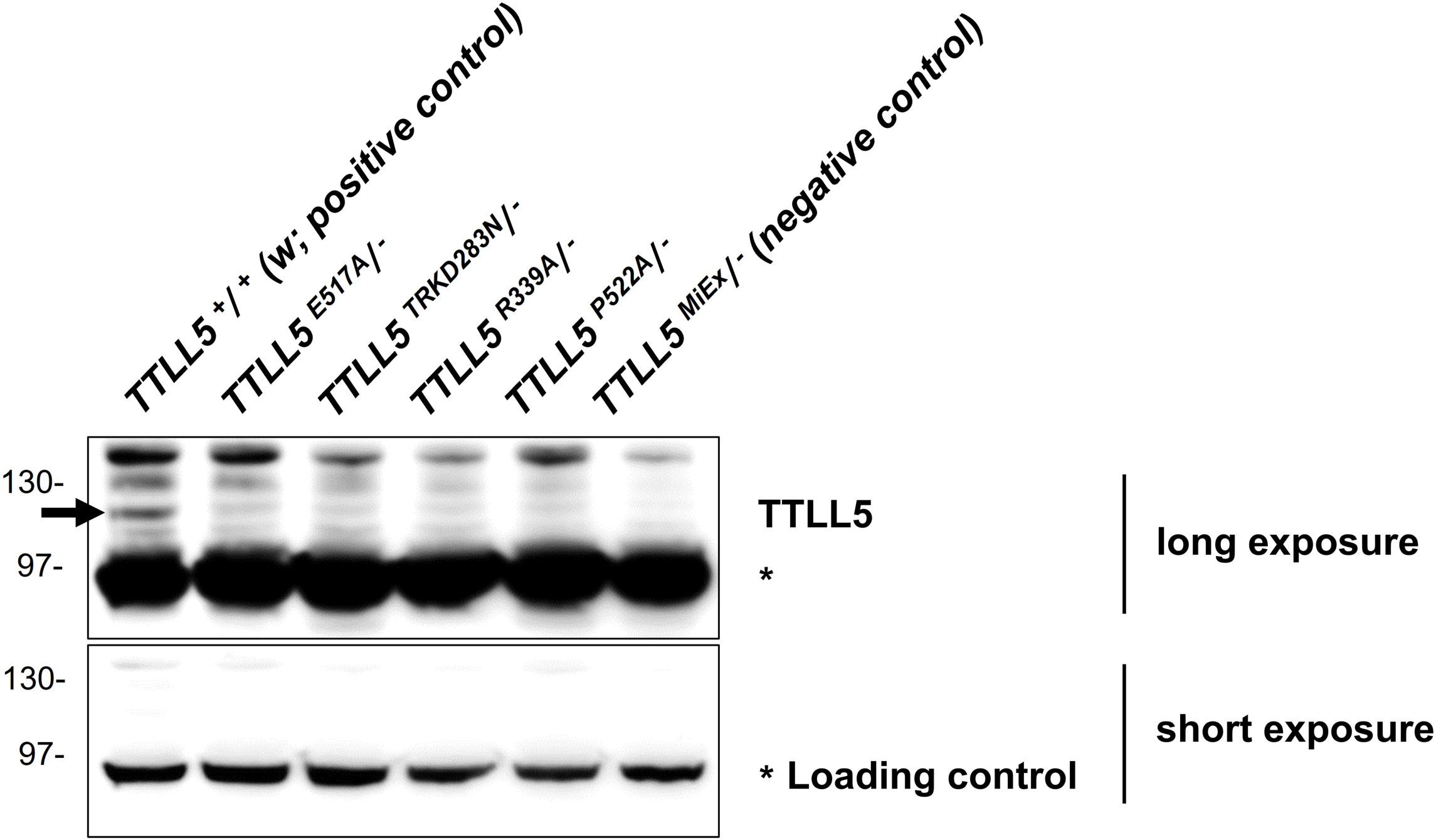
TTLL5 was stably expressed in all Crispr/Cas9 generated *TTLL5* mutants. All Crispr/Cas9 generated *TTLL5* mutations were analyzed hemizygously over *Df(3R)BSC679.* TR**K**D283N refers to a more complex mutation in which the 4 codons TR**K**D283 were replaced by a single N. We expect extracts from these animals to contain at most half the normal amount of TTLL5. *w* (*TTLL5^+^/TTLL5^+^*) was the positive control and the null mutation *TTLL5^MiEx^/Df(3R)BSC679* was the negative control. The signal of an unspecific band (labeled with *) under short exposure was treated as a loading control. *w,* which contains 2 copies of the *TTLL5^+^* allele expressed the highest levels of TTLL5. Crispr/Cas9 generated *TTLL5* mutants with a single allele of *TTLL5\** expressed less TTLL5 compared to *w,* but more compared to the null mutant *TTLL5^MiEx^/Df(3R)BSC679*. The TTLL5 levels of the hemizygous point mutants were similar to the ones of the hemizygous *TTLL5^+^* (see **Fig. 2B**). The faint band seen in *TTLL5^MiEx^/Df(3R)BSC679* is probably a background band.

**Fig. S2:**
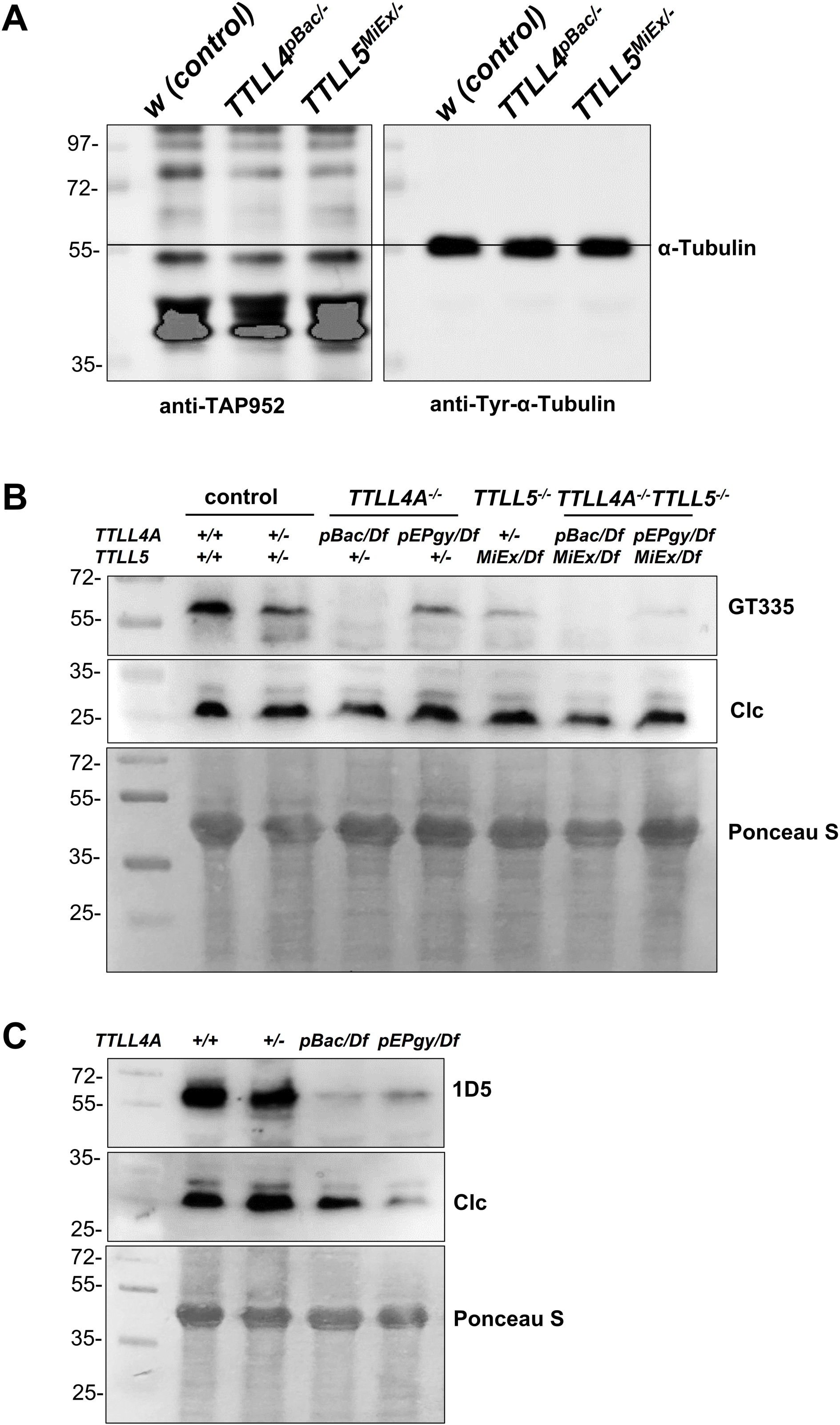
Preliminary studies on the role of *TTLL4* in α-tubulin modification in the *Drosophila* ovaries. A) The glycylation of α-tubulin was evaluated by Western blotting with the anti-monoglycylated Tubulin antibody TAP952. The anti-Tyr-α-Tubulin was a loading and size control. TAP952 did not detect a clear band in the α-Tubulin region in the wild type, *TTLL4A* or *TTLL5* mutant. B) Preliminary results indicate that *TTLL4A* is required for monoglutamylation and polyglutamylation of α-tubulin in ovaries. The monoglutamylation was evaluated by Western blotting with GT335 antibodies. The relative signal of GT335 is strongly reduced in *TTLL4A^pBac^, TTLL5^MiEx^, TTLL4A^pBac^;TTLL5^MiEx^, TTLL4A^pEPgy^;TTLL5 ^MiEx^* mutants compared to controls. GT335 signal is slightly reduced in *TTLL4A^pEPgy^* mutant. *TTLL4A^pEPgy^* is an insertion into the promoter region and appears to be a hypomorphic mutation while the *TTLL4A^pBac^* allele is an insertion into the leader and appears to be the stronger loss-of-function allele. Note that the lack of a GT335 signal in the *TTLL4A^-/-^*; *TTLL5^+/-^* is inconsistent with other results showing the role of *TTLL5* in monoglutamylation. Additional work is needed to find out what the reason for this is. C) The relative polyglutamylation signal visualized by the 1D5 antibodies is strongly reduced in the *TTLL4A^pBac^* mutant and also reduced in *TTLL4A^EPgy^*. Ponceau S and Clc were treated as loading controls. The *TTLL5* allele was analyzed in hemizygous animals over *Df(3R)BSC679*. All *TTLL4A* alleles were hemizygous over *Df(2L)BSC145.* The genotype for controls were *w* (+/+), and *+/Df(2L)BSC145 or TTLL4A^pBac^;+/ Df(3R)BSC679 or TTLL5 ^MiEx^*.

**Fig. S3:**
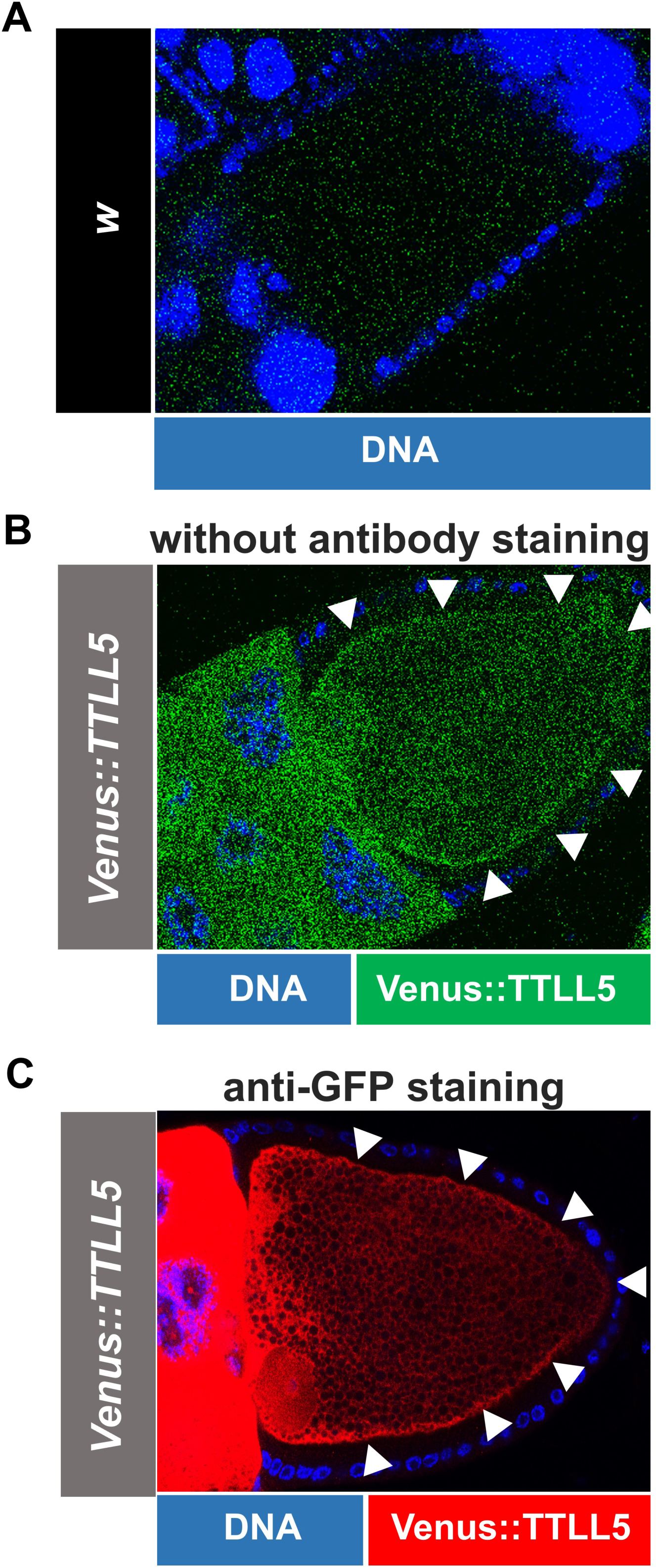
*Venus::TTLL5* expression in S10 oocytes. Confocal micrographs showing the S10 oocyte with or without anti-GFP staining. A) *w* oocyte shows the autofluorescence background of the oocyte under the microscope. B) The oocyte overexpressing *Venus:TTLL5* shows the live fluorescence of Venus (green). C) The oocyte overexpressing *Venus:TTLL5* shows the Venus::TTLL5 protein after staining with an anti-GFP antibody (red). Hochest was in blue. The cortical signal of Venus::TTLL5 in the oocytes is pointed out with white arrowheads. The genotype for *Venus:TTLL5* overexpression was *MattubGal4/UAS-Venus::TTLL5*.

**Fig. S4:**
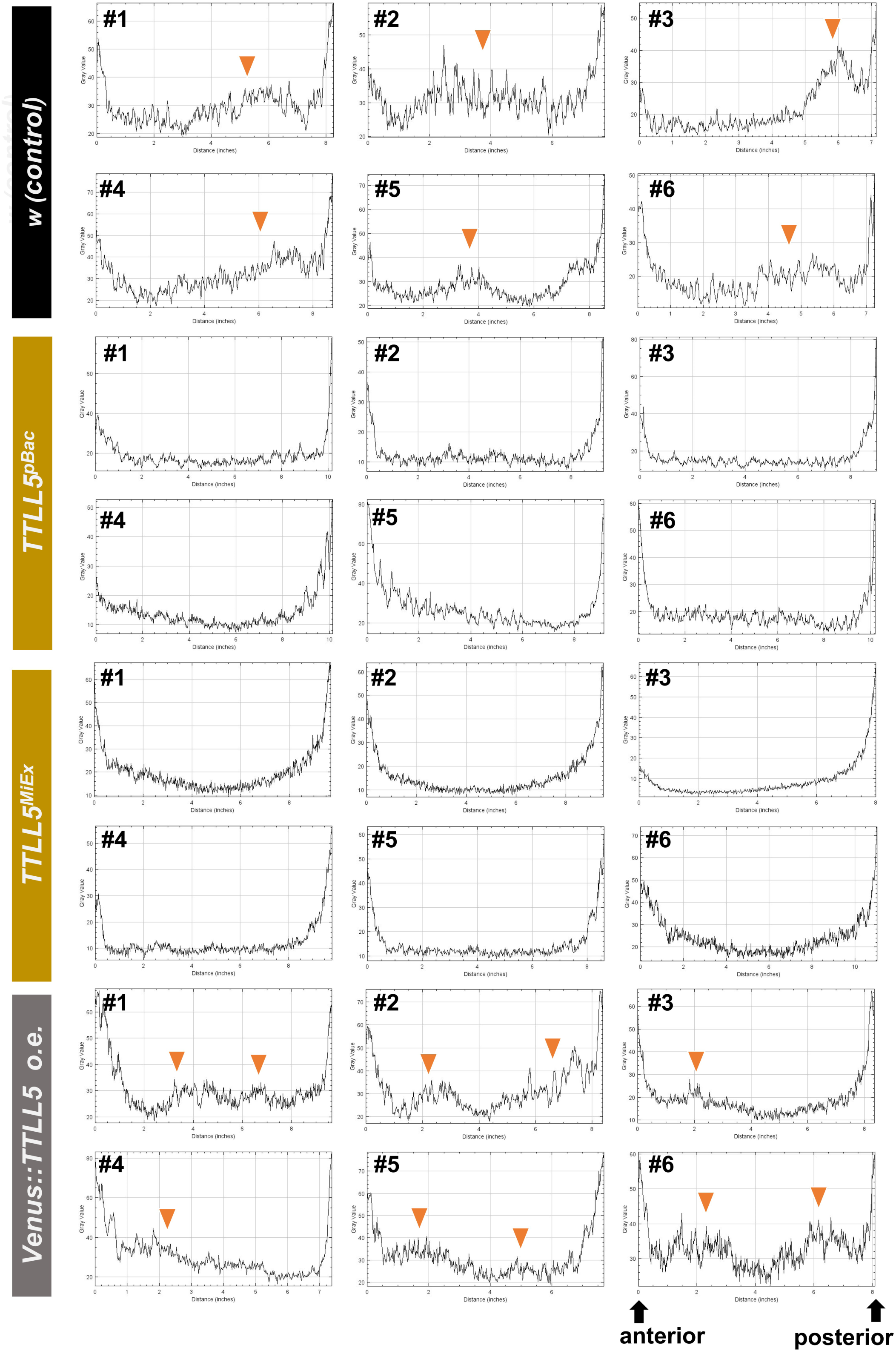
Khc distribution in S10B oocytes. Intensity charts from 6 samples of each genotype were plotted based on the line drawn along the AP axis of S10B oocytes (see Fig. 7E). The orange triangles in the control and *o.e. Venus::TTLL5* point out the inner regions accumulating higher levels of Khc. The control was *w* or *+/Df(3R)BSC679.* Genotypes for *TTLL5* mutants: *MattubGal4/+;TTLL5^pBac^/Df(3R)BSC679* and *TTLL5^MiEx^/Df(3R)BSC679*, respectively. Genotype for *TTLL5* overexpression: *MattubGal4/UAS-Venus::TTLL5*.

**Fig. S5:**
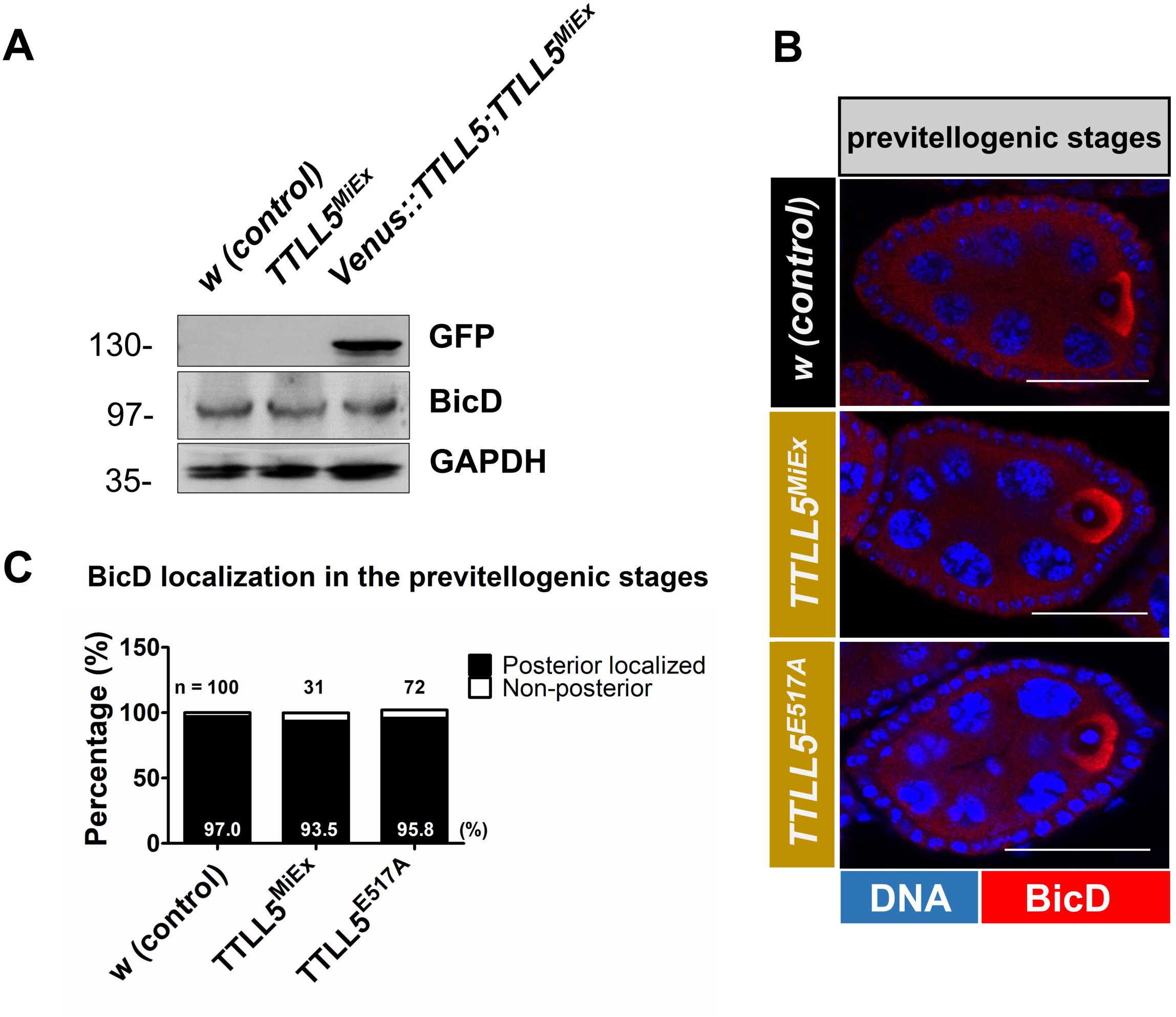
*TTLL5* has little or no effects on BicD expression and distribution in ovaries. A) BicD levels remained unchanged between the *w* control, a *TTLL5* deficiency mutant and the *TTLL5* rescue strain. GAPDH served as the loading control. B) Confocal micrographs show BicD localizing preferentially at the posterior of the previtellogenic oocytes of *w* controls and *TTLL5* mutants. Scale Bar: 25μm. BicD protein in red, and Hoechst in blue. C) 97%, 93.5% and 95.8% of the oocytes showed posteriorly localized BicD in the *w* control, *TTLL5^MiEx^* and *TTLL5^E517A^* mutants, respectively. The total numbers of early oocytes are indicated as n= in the chart. Genotype for control: *w* or *+/Df(3R)BSC679.* Genotypes for *TTLL5* mutants: *TTLL5^MiEx^/Df(3R)BSC679* and *TTLL5^E517A^/Df(3R)BSC679*, respectively.

**Fig. S6:**
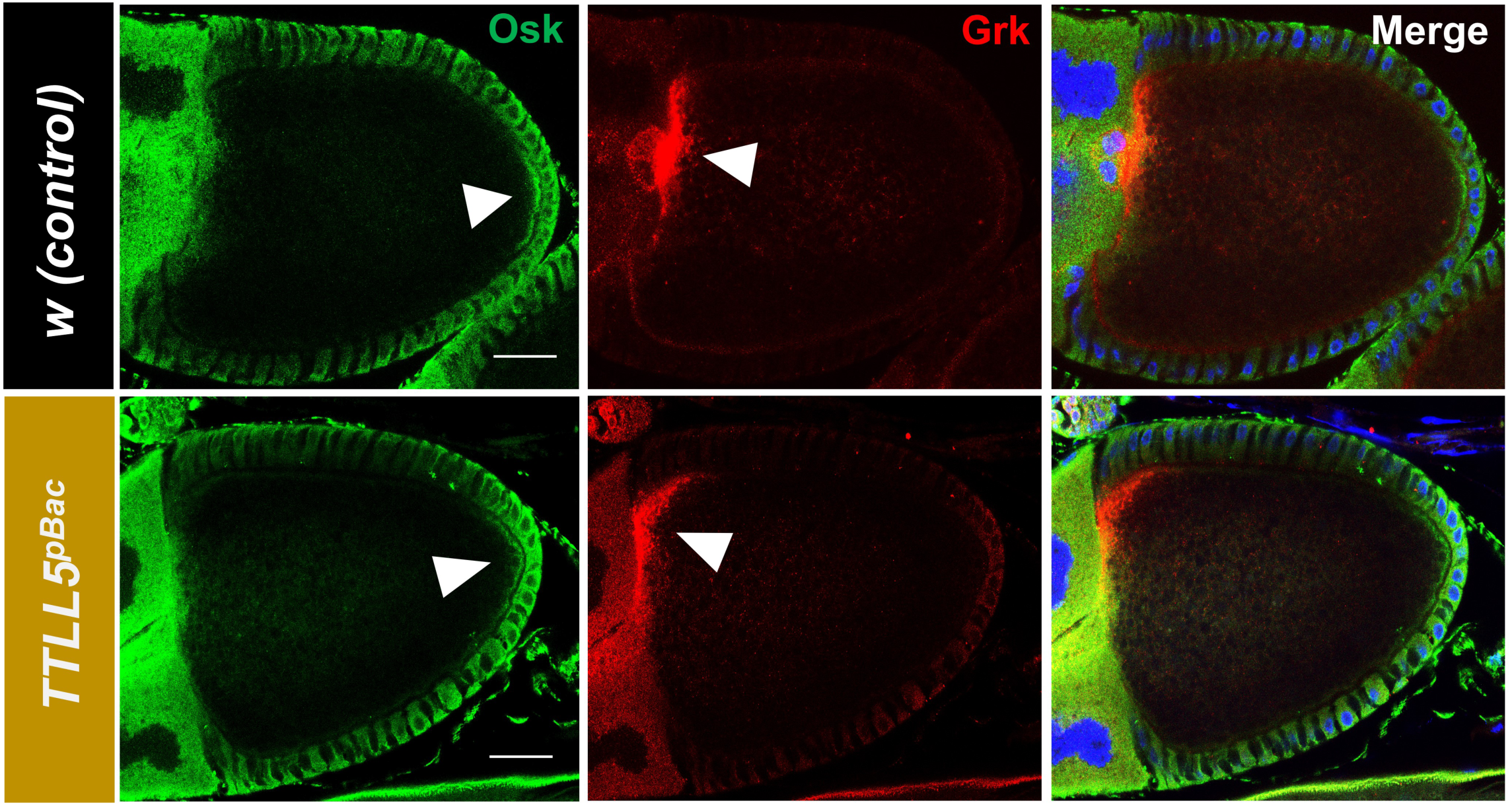
The polarity of S10B oocytes is normal in *TTLL5* mutant. In *TTLL5^pBac^* oocytes, Gurken (red) localizes properly to the dorsoanterior part of the oocyte cortex and Oskar (green) to the posterior cortex of 10B stage oocytes, indicating the polarity of MTs at this stage was normal. Blue channel: DNA. Genotypes for *TTLL5^pBac^*: *TTLL5^pBac^ /Df(3R)BSC679*.

**Mov. S1: Ooplasmic streaming in *w* (control; 2 oocytes)**

**Mov. S2: Ooplasmic streaming in *TTLL5^MiEx^* (2 oocytes)**

**Mov. S3: Ooplasmic streaming in *TTLL5^E517A^* (2 oocytes)**

**Mov. S4: Ooplasmic streaming in *Venus::TTLL5; TTLL5^MiEx^* (2 oocytes)**

**Mov. S5: Ooplasmic streaming in *Venus::TTLL5* (2 oocytes)**

**Mov. S6: Appl-Gal4>Mito::GFP trafficking in the L1 vein**. Top movie: control; bottom movie: *TTLL5^E517A^*.

## Notes

### Competing Interest Statement

The authors have declared no competing interest.

### Summary of Updates

This update contains minor improvements.

